# Systematic mapping of genetic interactions for *de novo* fatty acid synthesis

**DOI:** 10.1101/834721

**Authors:** Michael Aregger, Keith A. Lawson, Maximillian Billmann, Michael Costanzo, Amy H. Y. Tong, Katherine Chan, Mahfuzur Rahman, Kevin R. Brown, Catherine Ross, Matej Usaj, Lucy Nedyalkova, Olga Sizova, Andrea Habsid, Judy Pawling, Zhen-Yuan Lin, Hala Abdouni, Alexander Weiss, Patricia Mero, James W. Dennis, Anne-Claude Gingras, Chad L. Myers, Brenda J. Andrews, Charles Boone, Jason Moffat

**Affiliations:** Donnelly Centre, University of Toronto, 160 College Street, Toronto, Ontario, M5S 3E1, Canada; Department of Molecular Genetics, University of Toronto, 160 College Street, Toronto, Ontario, M5S 3E1, Canada; Division of Urology, Department of Surgery, University of Toronto; Department of Computer Science and Engineering, University of Minnesota – Twin Cities, 200 Union Street, Minneapolis, MN, 55455, USA; Lunenfeld-Tanenbaum Research Institute, Mount Sinai Hospital, Toronto, Ontario, M5G 1X5, Canada; Bioinformatics and Computational Biology Graduate Program, University of Minnesota – Twin Cities, 200 Union Street, Minneapolis, MN, 55455, USA; Institute for Biomaterials and Biomedical Engineering, 160 College Street, Toronto, Ontario, M5S 3E1, Canada

## Abstract

The *de novo* synthesis of fatty acids has emerged as a therapeutic target for various diseases including cancer. While several translational efforts have focused on direct perturbation of *de novo* fatty acid synthesis, only modest responses have been associated with mono-therapies. Since cancer cells are intrinsically buffered to combat metabolic stress, cells may adapt to loss of *de novo* fatty acid biosynthesis. To explore cellular response to defects in fatty acid synthesis, we used pooled genome-wide CRISPR screens to systematically map genetic interactions (GIs) in human HAP1 cells carrying a loss-of-function mutation in *FASN*, which catalyzes the formation of long-chain fatty acids. *FASN* mutant cells showed a strong dependence on lipid uptake that was reflected by negative GIs with genes involved in the LDL receptor pathway, vesicle trafficking, and protein glycosylation. Further support for these functional relationships was derived from additional GI screens in query cell lines deficient for other genes involved in lipid metabolism, including *LDLR*, *SREBF1*, *SREBF2*, *ACACA*. Our GI profiles identified a potential role for a previously uncharacterized gene *LUR1* (*C12orf49*) in exogenous lipid uptake regulation. Overall, our data highlights the genetic determinants underlying the cellular adaptation associated with loss of de novo fatty acid synthesis and demonstrate the power of systematic GI mapping for uncovering metabolic buffering mechanisms in human cells.

## INTRODUCTION

Lipid metabolism as a source of energy for cancer cells, supporting rapid cell division and contributing to cell survival, and fatty acid derivatives play key roles in oncogenic signalling. Alterations in lipid metabolism, specifically the uptake of lipids and/or synthesis of fatty acids, comprise different aspects of metabolic reprogramming that are well documented in cancer and other indications, including metabolic syndrome and fatty liver disease (Chen & Huang, 2019). *De novo* fatty acid synthesis has gained significant traction as a targetable pathway following observations that overexpression of *FASN,* which encodes fatty acid synthase and catalyzes the formation of long chain fatty acids, and *ACACA*, which codes for Acetyl-CoA Carboxylase Alpha and acts directly upstream of FASN, are associated with decreased survival rates for numerous solid malignancies (Chen *et al*, 2019; Imoto, 2018; Menendez & Lupu, 2017; Garber, 2016; Röhrig & Schulze, 2016). Efforts to develop and translate small molecule inhibitors of FASN (e.g. TVB-2640) have helped validate this enzyme as a targetable liability in cancer (Jones & Infante, 2015; Benjamin *et al*, 2015), and have led to several clinical trials (e.g. *NCT02223247*, *NCT02948569*, *NCT03179904*, *NCT02980029*). Given that metabolic pathways are highly buffered to deal with environmental change, genetic screening approaches are a powerful strategy to reveal metabolic regulatory mechanisms that underscore metabolic redundancy, cross-talk and plasticity (Birsoy *et al*, 2014, 2015). An understanding of how cells adapt to perturbation of *de novo* fatty acid synthesis could help identify new targetable vulnerabilities that may inform novel therapeutic strategies or biomarker approaches.

Mapping genetic interaction (GI) networks provides a powerful approach for identifying the functional relationships between genes and their corresponding pathways. The systematic exploration of pairwise GIs in model organisms revealed that GIs often occur among functionally related genes and that GI profiles organize a hierarchy of functional modules (Costanzo *et al*, 2016; Fischer *et al*, 2015). Thus, GI mapping has become an effective strategy for identifying functional modules and annotating the roles of previously uncharacterized genes. Model organism GI mapping has also provided insight into the mechanistic basis of cellular plasticity or phenotypic switching that occurs as cells evolve within their environments (Harrison *et al*, 2007; Szappanos *et al*, 2011). Accordingly, the insights gained through systematic interrogation of GIs have fuelled significant interest to leverage these approaches towards functionally annotating the human genome.

Recent technological advances using CRISPR-Cas enable the systematic mapping of GIs in human cells (Wright *et al*, 2016; Doench, 2018). Here, we explore genome-wide GI screens within the context of human query mutant cells defective for *de novo* fatty acid synthesis. We systematically mapped genome-wide GI profiles for six genes involved in lipid metabolism, revealing cellular processes that pinpoint genetic vulnerabilities associated with defects in *de novo* fatty acid synthesis. In particular, negative GIs with known fatty acid synthesis genes tend to identify other genes that are associated with this process, including a previously uncharacterized gene *C12orf49* (*LUR1*), which appears to function as a regulator of exogenous lipid uptake. Collectively, our data support the strategy of systematically mapping digenic interactions using knockout query cell lines for identifying buffering mechanisms within metabolism.

## RESULTS

### Systematic identification of genetic interactions for *de novo* fatty acid synthesis

*De novo* fatty acid synthesis is a multi-step enzymatic process that converts cytosolic acetyl-CoA, malonyl-CoA, and NADPH to palmitate. Palmitate can be used directly or further elongated and/or undergo desaturation to form alternate lipid species. To systematically identify GIs associated with this metabolic process, we performed genome-wide CRISPR screens in coisogenic cell lines either wild-type or deficient in FASN, a *de novo* fatty acid synthesis enzyme that is frequently overexpressed in malignancies (Röhrig & Schulze, 2016; Currie *et al*, 2013) (Figure 1a). We chose the human near-haploid cell line HAP1 as a model system, given the relative ease for generating knockout (KO) mutations in this background (Carette *et al*, 2011). We first validated our clonal *FASN*-KO cells by confirming loss of FASN protein levels by western blot (Figure S1a). We also performed targeted metabolite profiling of our parental HAP1 and *FASN*-KO cells, which revealed a significant increase in the FASN substrate malonyl-CoA in the *FASN*-KO cells, demonstrating their suitability as a model system for defective *de novo* fatty acid synthesis (Figure S1b).

**Figure 1.**
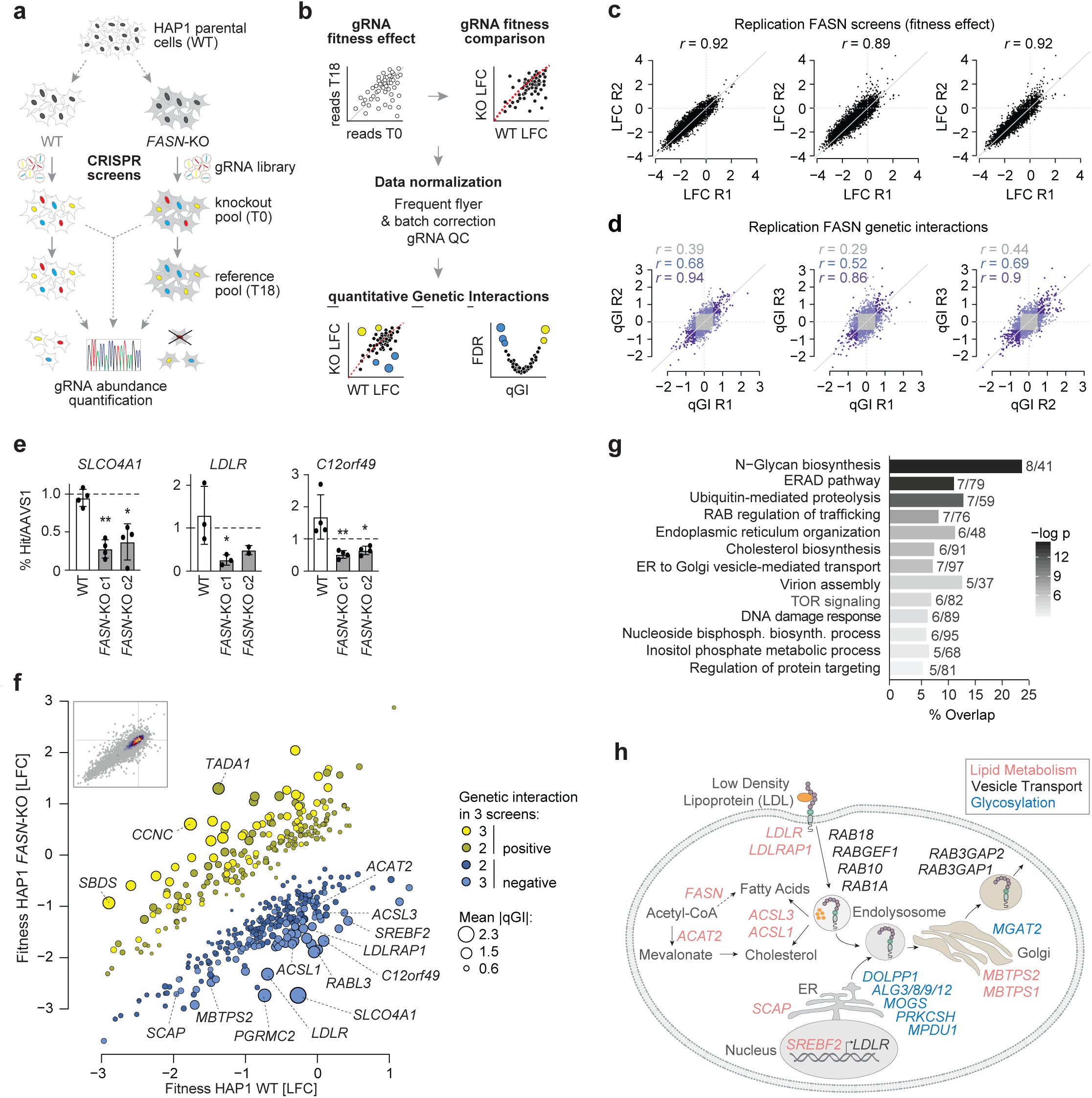
Genome-scale identification of digenic interactions with *FASN*. (**a**) Schematic outline for the identification of genetic interactions in coisogenic HAP1 cell lines. *FASN* knockout (KO) and wildtype parental cells are infected with a lentiviral genome-wide CRISPR gene KO library (TKOv3) and gRNA abundance is determined using Illumina sequencing of guide RNA (gRNA) sequences amplified from extracted genomic DNA from the starting cell population (T0) and end time point (day 18, T18) of the screen. (**b**) Schematic outline for scoring quantitative genetic interactions (qGI) across coisogenic query cell lines. First, the log_2_ fold-change (LFC) for each gRNA comparing sequence abundance at the starting (T0) and end time point (T18) in a given query KO or wildtype (WT) cell population are computed. Differential LFC for each gRNA are then estimated by comparing its LFC in WT and query KO cells. A series of normalization steps and statistical tests are applied to these data to generate gene-level qGI scores and false discovery rates (see Methods for details). The LFC scatterplot (bottom left graph) visualizes differential fitness defects in a specific query KO and WT cells, whereas the volcano plot (bottom right graph) visualizes qGI scores for a specific query. (**c**) Replicate analysis of gene loss of function fitness phenotypes in FASN screens. Scatter plots of LFC associated with perturbation of 17,804 individual genes derived from a *FASN* query *KO* mutant screen conducted in triplicate. Reproducibility of fitness effects were determined by measuring Pearson correlation coefficients (r) between all possible pairwise combinations of *FASN*-KO replicate screens. (**d**) Evaluation of *FASN* quantitative genetic interactions (qGIs). qGI scores were measured by comparing the LFC for every gene represented in the TKOv3 gRNA library in a *FASN*-KO with those observed in a WT cell line, as described. Scatter plots show *FASN* genetic interactions (qGI scores) derived from all possible pairwise combinations of three biological replicate screens. The Pearson correlation coefficient (*r),* based on comparison of all qGI scores (r shown in grey, calculated on all the grey, blue and purple data points in the scatter plots), or only genetic interactions that exceed a given significance threshold (|qGI| > 0.5, FDR < 0.5) in one (blue) or two screens (purple). (**e**) Validation of FASN negative genetic interaction. Bar plots depict the ratio of WT and *FASN*-KO (2 independent clones, c1 and c2) cells carrying a gRNA targeting *SLCO4A1, LDLR* or *C12orf49,* which all showed a negative interaction with *FASN,* compared to a gRNA targeting *AAVS1* (intergenic control). Experiments were performed with three independent gRNAs targeting each genetic interaction screen hit. All data are represented as means ± standard deviation (n = 3 or 4), **p < 0.01, and *p < 0.05; one-way ANOVA. (**f**) *FASN* negative and positive genetic interactions. A scatter plot illustrating the fitness (LFC) of 450 genes individually targeted in a *FASN*-KO vs. WT parental HAP1 cell line. Each of the 450 genes shown exhibited a significant genetic interaction in at least 2 out of 3 *FASN*-KO replicate screens (|qGI| > 0.5, FDR < 0.5). Negative (blue) and positive (yellow) *FASN* genetic interactions are shown. Node size corresponds to the mean absolute qGI score derived from 3 replicate screens. Genes with mean absolute qGI score > 1.5 as well as selected negative interactions involving genes with established roles in lipid metabolism are indicated. [Inset] Scatter plot of FASN genetic interaction scores for all 17,804 genes targeted by the TKOv3 gRNA library where the color indicates density of genes. (**g**) Enrichment for Gene Ontology (GO) molecular function, GO bioprocesses and Reactome terms among genes that exhibited a significant negative genetic interaction with *FASN* (significant in at least two *FASN* replicates, |qGI| > 0.5, FDR < 0.5). The number of genes annotated to each term and shown to interact with FASN are indicated. (**h**) Schematic depicting the function of selected *FASN* negative interactions known to be involved in lipid uptake and homeostasis pathways (red), vesicle transport (black) and glycosylation (blue).

To map *FASN* GIs, we performed genome-wide CRISPR screens using the sequence optimized TKOv3 gRNA library (Hart *et al*, 2017) in both the *FASN*-KO query cell line and control wild-type (WT) HAP1 cells, and we compared the relative abundance of individual gRNAs between the screen start (T0) and end (T18) time points (Figure 1a-b). The relative abundance of gRNAs targeting each of ∼18,000 genes in WT cells provides an estimate of single mutant fitness, whereas the relative abundance of gRNAs in a query mutant cell line provides an estimate of double mutant fitness. Since mutant phenotypes can strongly depend on culture conditions (Billmann *et al*, 2018) and most standard cell culture media contains supra-physiological nutrient levels that could mask phenotypic effects of perturbing certain metabolic pathways, we performed our screens utilizing media conditions containing the minimum amounts of glucose and glutamine required to sustain proliferation of HAP1 cells; termed limiting media (Figure S1c, see Methods).

We developed a quantitative genetic interaction (qGI) score that measures the strength and significance of genetic interactions by comparing the relative abundance of gRNAs in a given query mutant cell line to the relative abundance of gRNAs targeting the corresponding genes in an extensive panel of 21 genome-wide WT HAP1 screens (Figure 1b, see Methods). In this context, negative interactions are identified as genes whose corresponding gRNAs exhibit significantly decreased abundance in a mutant KO background relative to the control WT HAP1 cell line, whereas positive interactions reflect genes with increased gRNA abundance in a mutant cell line relative to the parental line.

We performed three independent genome-wide, GI screens using our *FASN*-KO query mutant cell line. Because GIs rely on accurate measurement of single and double mutant phenotypes, we first examined the reproducibility of our single and double mutant fitness measurements (see Methods). We observed a strong agreement of single gene fitness effects (LFC) among 21 replicate WT HAP1 (r > 0.87) (Figure S1d) and double mutant fitness effects derived from independent *FASN*-KO replicate screens (r > 0.89) (Figure 1c). Moreover, all three *FASN* screens robustly discriminated a set of reference essential genes from non-essential genes (Figures S1e-f).

The identification of qGI scores depends on comparison of single mutant fitness measurements in a WT HAP1 cell screen and double mutant fitness measurements in a query mutant screen, both of which have inherent variability associated with them; therefore, the reproducibility of qGIs is expected to be more challenging than the measurement of either single or double mutant fitness phenotypes. Indeed, modest agreement was observed between qGI scores of the three *FASN*-KO replicate screens prior to filtering for significant interactions (pairwise r = 0.29 to 0.44) (Figure 1d). The pairwise correlation between replicate screens increased substantially when we considered GIs found to be significant (|qGI| > 0.5, FDR < 0.5) in at least one (r = 0.52-0.69) or two (r = 0.86-0.94) *FASN*-KO replicate screens (Figure 1d, **Table S1**).

Leveraging all 3 *FASN-KO* replicates, we developed a reproducibility score that measures each gene’s contribution to the covariance within two replicate screens and summarizes the resulting values across all available screen pairs (replicate 1-2, 1-3, 2-3) (Methods, **Table S1**). This analysis confirms that both the strongest positive and negative qGI scores were highly reproducible across independent screens (Figure S1g). In particular, the most reproducible negative GIs with *FASN* were interactions with *SLCO4A1*, *PGRMC2*, *LDLR, RABL3* and *C12orf49* (Figure S1g, **Table S1**). We tested three of these top five strongest negative GIs by independent validation assays and confirmed all three, examining WT and *FASN*-KO HAP1 cells expressing gRNAs against *SLCO4A1, LDLR* and *C12orf49* (Figure 1e, S1h).

To generate an aggregate set of FASN GIs, we mean-summarised qGI scores across the three replicate screens (Figure 1f, **Table S2**). At a pathway level, significant negative GIs (qGI < −0.5, FDR < 0.5) with *FASN* were strongly enriched for genes annotated with roles in protein glycosylation, vesicle transport and cholesterol metabolism (FDR <0.05) (Figure 1g, **Table S3**). In the global yeast genetic network negative GIs often connect functionally related genes (Costanzo *et al*, 2010, 2016), and we observed a similar general trend for the *FASN* negative GIs. For example, the *FASN* negative GIs included genes with established roles in the uptake, transport, and breakdown of low density lipoprotein (LDL), a major extracellular source of lipids, including the LDL receptor (*LDLR*) itself and its coreceptor adaptor protein (*LDLRAP1*). We also observed negative GIs between *FASN* and the transcription factor *SREBF2*, which controls expression of *LDLR*, as well as *SCAP*, *MBTPS1* and *MBTPS2*, all of which are important for the activation and nuclear translocation of SREBF2 upon cholesterol depletion (Figure 1h). Moreover, we observed negative GIs with additional lipid metabolic processes such as cholesterol biosynthesis (*ACAT2*), genes functioning in long chain fatty acid activation and β-oxidation (*ACSL1, ACSL3*), and vesicle trafficking genes (*RAB18/10/1A, RABGEF1, RAB3GAP2/1*) (Figures 1h, S1i), as well as a positive GI with the gene encoding stearoyl-CoA desaturase (*SCD*), the product of which catalyses the rate-limiting step in the biosynthesis of monounsaturated fatty acids.

The *FASN* screen also highlighted an enrichment for genes functioning in protein N-linked glycosylation (e.g. *ALG3/8/9/12, MOGS, DOLPP1, PRKCSH, MGAT2*) (Figures 1g-h, S1i). Interestingly, the hexosamine biosynthetic and N-linked glycosylation pathways have been implicated in facilitating lipid accumulation from environmental sources through direct modulation of N-glycan branching on fatty acid transporters, possibly explaining the strong GIs we observe (Ryczko *et al*, 2016). N-linked glycosylation is also known to play an important role in the activity of LDLR and activation of the SREBP transcriptional programs, providing a potential explanation for the interaction between loss of *FASN* and the glycosylation pathway (Cheng *et al*, 2015; Wang *et al*, 2018). Finally, we observed a significant negative GI between *FASN* and *SLCO4A1* (Figures 1f, S1g). *SLCO4A1* encodes a member of the organic anion-transporting polypeptides (OATPs), which can transport a wide range of structurally unrelated compounds including hormones, bile acids and lipid species (prostaglandins) (Obaidat *et al*, 2012). To summarize, these results suggest that in the absence of cell autonomous *de novo* fatty acid synthesis, cells depend on uptake and breakdown of lipids from the environment or the synthesis of sterols, with our data illuminating the genetic determinants of how cells rewire to meet the demand for lipids in proliferating cells.

### Expanding the genetic interaction landscape of *de novo* fatty acid synthesis

To better understand the GI landscape of *de novo* fatty acid synthesis, we next performed pooled genome-wide CRISPR screens using the TKOv3 library in five additional coisogenic cell lines harbouring genetic KO of genes that exhibited significant negative GIs with our *FASN*-KO query, including *LDLR*, *C12orf49* and *SREBF2* (**Table S2**), as well as two genes that did not show a negative GI with *FASN*, including *SREBF1*, which regulates the expression of *FASN* and other *de novo* fatty acid genes, *and ACACA,* which functions in the same pathway and immediately upstream of *FASN* (Figure 2a) (Röhrig & Schulze, 2016; Currie *et al*, 2013; Horton *et al*, 2008). Each of these five query gene screens was performed in technical triplicate (i.e. parallel cultures from a common infection). Since these additional GI screens were performed under the same conditions as we used for the *FASN*-KO screens, we applied the same confidence threshold on the derived qGI scores (|qGI| > 0.5, FDR < 0.5; Methods) (Figures 2b-f, S2a-b, **Table S2**). At this confidence threshold, we estimated a per-screen false discovery rate of ∼0.3 and a false negative rate of ∼0.6 (Methods; Figure S1j).

**Figure 2.**
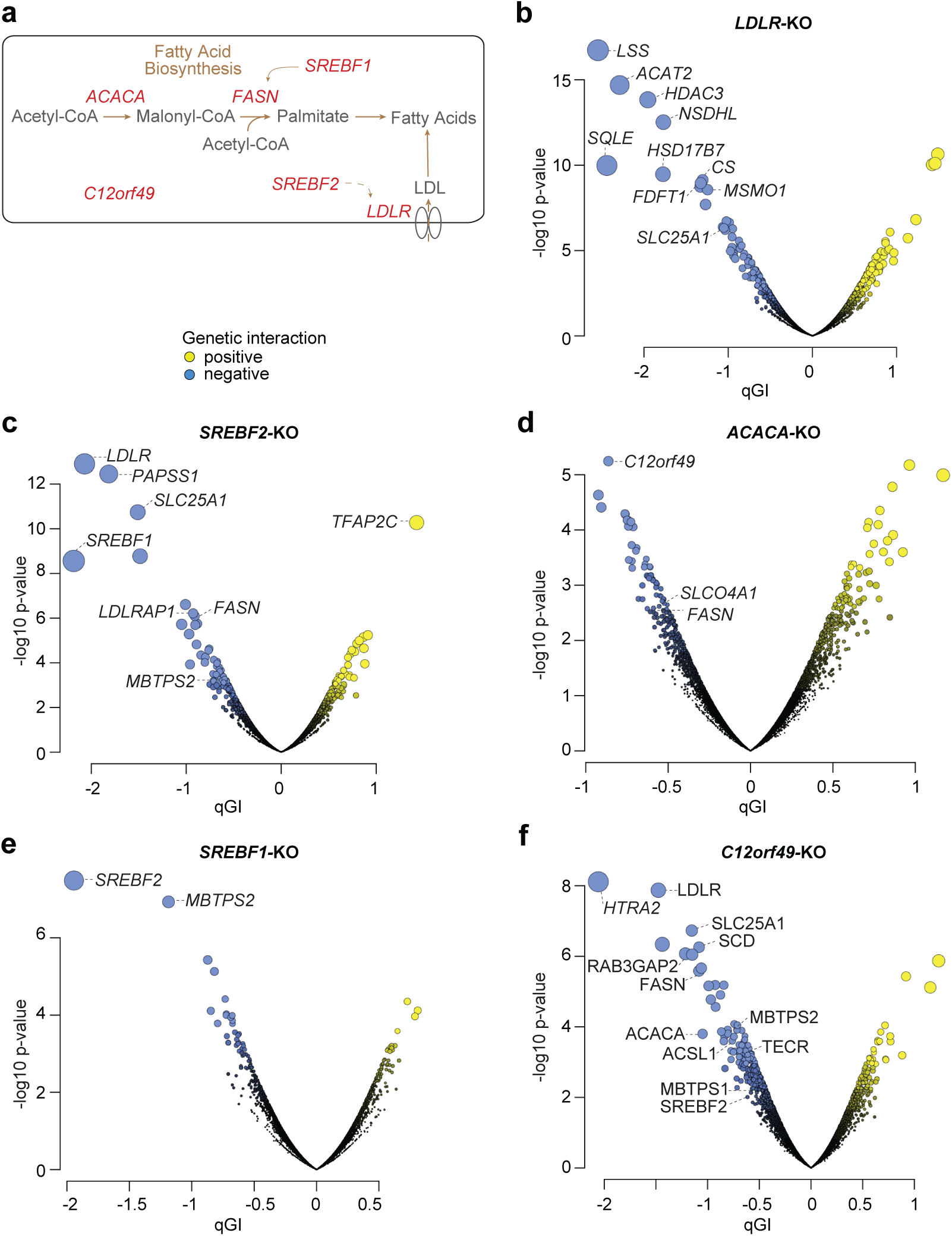
Querying five additional lipid metabolism genes for digenic interactions. (**a**) Schematic diagram showing key steps in fatty acid metabolism. The genes encoding the proteins mediating these key steps, which are also query genes for genetic interaction screens described in the main text, are labelled in red. (**b-f**) Volcano plots showing qGI scores versus false discovery rates (-log10 p-value) for the results of the (b) *LDLR*-KO, (c) *SREBF2*-KO, (d) *ACACA*-KO, (e) *SREBF1*-KO and (f) *C12orf49*-KO screens. Colored dots indicate genes that meet the standard threshold of |qGI| > 0.5, FDR < 0.5, where positive GIs are indicated in yellow and negative GIs in blue. The dot size is proportional to both qGI and FDR, calculated as described in the methods. Genes with |qGI| scores > 1.5 as well as selected top negative GI hits associated with lipid metabolism, citrate synthesis and transport are indicated.

We next analyzed the functional enrichment across all GIs identified by our fatty acid synthesis-related query screens. While the positive GIs were not functionally informative in general, we observed a clear 5-fold enrichment of negative GIs for genes annotated to functionally relevant pathways, which were defined by the metabolism-focused HumanCyc standard (Figure S2c) (Romero *et al*, 2005). We further quantified enrichment for pathways annotated at different levels of the HumanCyc database hierarchy, including gene sets corresponding to general metabolic reaction categories, sub-categories, and finally specific metabolic pathways (**Table S4**). At the most general level of the HumanCyc pathway hierarchy, negative GIs from all six genome-wide screens were most enriched for genes annotated to the biosynthesis and macromolecule modifications pathway categories (Figure 3a). Further analysis of these terms at a more specific level of the HumanCyc hierarchy (i.e. sub-category level), we found that genes exhibiting negative GIs were associated with functions related to the roles of our six query genes, including fatty acid, lipid and carbohydrate biosynthesis (Figures 3b, S3a). At a more refined level of functional specificity within the fatty acid and lipid biosynthesis pathway, we found that each query gene was associated with a significant enrichment for negative GIs with functionally-related genes of distinct pathways. For example, the *LDLR* GI profile includes negative GIs with genes in the cholesterol/epoxysqualene biosynthesis pathway (i.e. *HMGCS1*, *MSMO1*, *HMGCR, FDFT1, NSDHL, HSD17B7, SQLE, HSD17B7, ACT2, SQLE, LSS*) and the *ACACA*, *LDLR* and *SREBF2* GI profiles include negative GIs with fatty acid elongation and biosynthesis pathway genes (*FASN, ACACA, OXSM*) (Figures 3c-d). Notably, the *FASN* GI profile, and to a lesser extent the *ACACA* and *LDLR* GI profiles, revealed negative GIs with pathways and genes involved in N-glycosylation initiation (*ALG6*, *ALG13*, *ALG11*, *ALG1*, *ALG2*, *ALG8*, *ALG5*, *ALG3*, *ALG12*, *ALG9*), processing (*MOGS*, *PRKCSH*), dolichol monophosphate mannose synthase activity (*DPM2*, *DPM3*, *DPM1*), and glycan transfer (*STT3A*, *STT3B*) (Figures 3c,e, **Table S4**).

**Figure 3.**
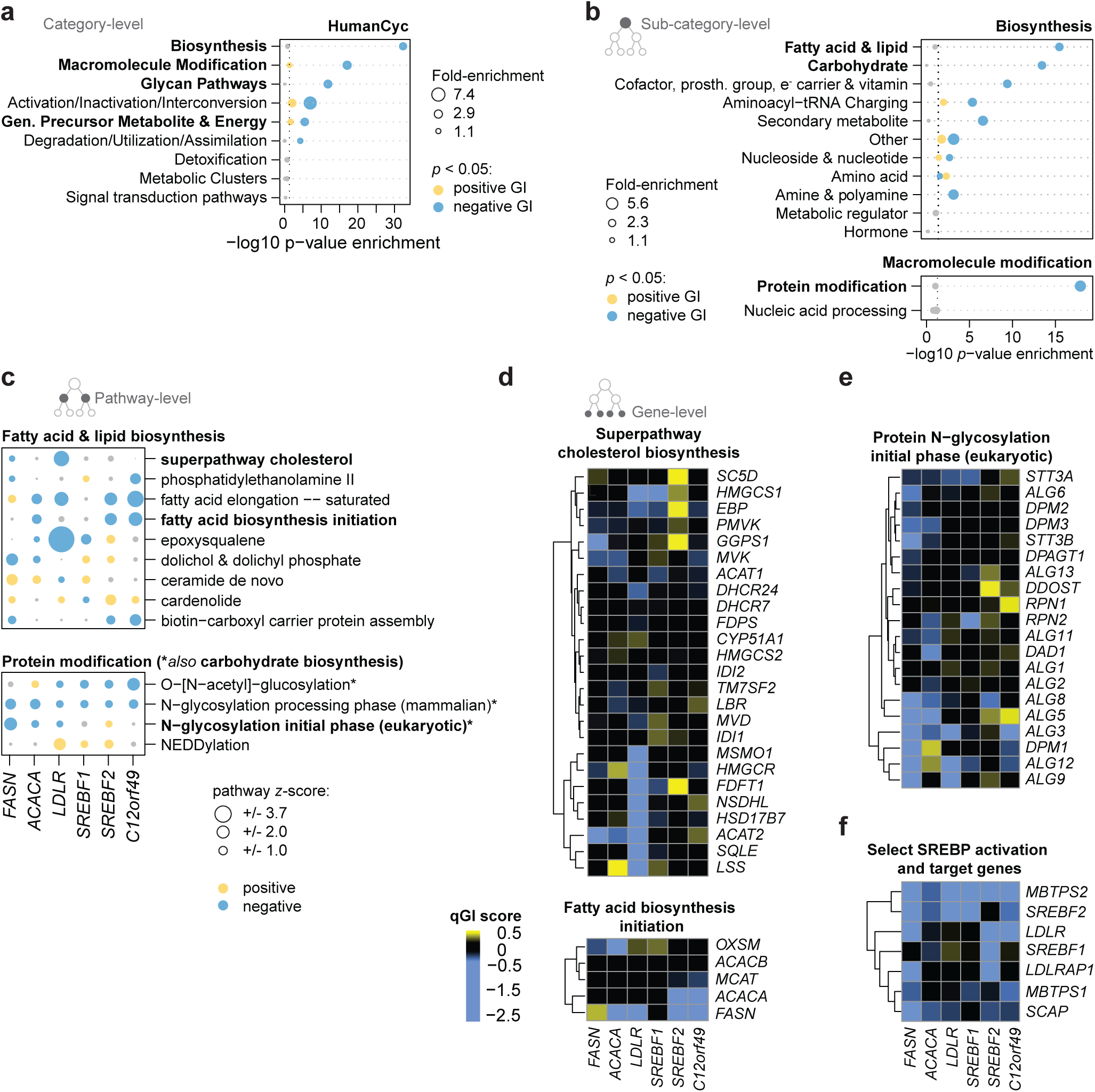
Genetic interactions reveal multiple levels of functional enrichment. (**a**) Dot plot of normalized pathway enrichment scores on the HumanCyc category level, calculated from qGIs across all six query genes (*FASN, C12orf49, LDLR, SREBF2, ACACA, SREBF1*). A GI is identified for a query-library pair if the |qGI| > 0.5 and FDR < 0.5. Enrichment for positive (yellow) and negative (blue) GIs is tested in each of the 10 HumanCyc main pathway categories using a hypergeometric test. Enrichment with p-value < 0.05 are blue (negative GI) and yellow (positive GI). Dot size is proportional to the fold-enrichment in the indicated categories and specified in the legend. Categories indicated in bold are further expanded in part (b) and in Supplemental Figure 3a. (**b**) Dot plot of normalized pathway enrichment of GIs on a sub-category level, calculated as described in part (a), except that sub-categories were examined inside the Biosynthesis and Macromolecule Modification HumanCyc branches. Enrichment with p-value < 0.05 are blue (negative GI) and yellow (positive GI). Dot size is proportional to the fold-enrichment in the indicated categories and specified in the legend. Categories indicated in bold text are further expanded in part (c). (**c**) Matrix dot plot of pathway enrichments of GIs for the fatty acid and lipid biosynthesis and protein modification sub-categories. Dots show positive (yellow) or negative (blue) z-transformed qGI scores summarized at a pathway-level. qGI scores were first z-score transformed at a gene-level for each genome-wide query screen separately. Then, a mean z-score was calculated for each pathway for a given query screen. Dot size corresponds to the absolute z-transformed mean qGI score, grey dots represent |z| < 0.5. Pathways marked with an asterisk are annotated to both protein modification and carbohydrate biosynthesis pathways. Bold pathways are shown in (d-e). Pathways were displayed if they shared an absolute z-score larger than 1.5 with any query gene. (**d-f**) Gene-level heatmaps for genes involved in enriched pathways. qGI scores between query genes and all genes from the selected pathways. Positive and negative qGI scores are indicated in yellow and blue, respectively.

Our survey of GIs related to perturbation of *de novo* fatty acid synthesis or exogenous fatty acid uptake pathways provided unique insight into the genetic regulation of these processes. Specifically, for the *SREBF2* screen, while we observed negative GIs with lipid uptake genes such as *LDLR* and *LDLRAP1* (Figure 3f, **Table S2**), none were observed with the cholesterol biosynthesis pathway (Figures 3d, 2e). This observation is consistent with *SREBF2* being the predominant transcriptional regulator of cholesterol homeostasis (Horton *et al*, 2008); its perturbation does not further reduce cellular fitness in cells deficient for cholesterol biosynthesis. In addition, we also detected a strong positive GI between *SREBF2* and *TFAP2C* (Figure 2c). Indeed, the*TFAP2* transcription factor family has recently been proposed as a ‘master’ regulator of lipid droplet biogenesis (Scott *et al*, 2018), with our data suggesting that reduced sequestration of lipids into lipid droplets may benefit *SREBF2*-KO cells to mitigate lipid starvation.

In contrast*, SREBF1* did not show enrichment for GIs for either the cholesterol or fatty acid synthesis pathways (Figure 3c, **Table S2**). Instead, this query was found to show only a strong reciprocal negative GI with its paralog *SREBF2,* highlighting the functional redundancy between the paralog pair (Figure 2e, **Table S2**) and suggesting that *SREBF2* may regulate some of the transcriptional targets of *SREBF1* as previously described previously (Shimano & Sato, 2017; Horton *et al.,* 2008). Furthermore, the imbalanced number of GIs between *SREBF1* and *SREBF2* may point towards asymmetric paralog evolution, whereby duplicated genes gain or lose functional roles at different rates while maintaining partially redundant functions, a process previously observed in yeast and human cells (Zhou *et al*, 2014; VanderSluis *et al*, 2010; Ascencio *et al*, 2017).

### A novel role for *C12orf49* in lipid biosynthesis

One of the strongest negative GIs identified in both the *FASN* and the *ACACA* profiles involved the uncharacterized gene *C12orf49,* suggesting that this gene may have a role in lipid metabolism (Figures 1f, 2d, **Table S2**). C12orf49 is a 23.5 kDa protein that is part of the UPF0454 family of uncharacterized proteins, contains an N-terminal transmembrane sequence, single uncharacterized DUF2054 domain of approximately 200 amino acid residues, 14 conserved cysteines three of which are annotated to form CC-dimers, and a predicted glycosylation site (The UniProt Consortium, 2019)(Figure S4a). In some plant proteins, the uncharacterized UPF0454 is found situated next to a glycosyltransferase domain and thus may be targeted into the lumen of the ER or Golgi (Mitchell *et al*, 2019). By extension, the bulk of the C12orf49 protein may reside in the lumen of the ER or Golgi. In addition, C12orf49 is ubiquitously expressed across tissues and cell lines (http://www.proteinatlas.org) (Uhlen *et al*, 2015)). Notably, expression of *C12orf49* is associated with differential prognoses on univariate analysis of TCGA data across multiple tumor types, including kidney, breast, liver and sarcoma (Figures S4b-e; p < 0.05) (Nagy *et al*, 2018), which further motivated us to study the functional role of this previously uncharacterized gene.

Genetic interactions derived from a genome-wide screen using a *C12orf49*-KO query cell line further supported a role for this gene in lipid biogenesis. Consistent with the results described above, *C12orf49* showed a strong negative GI with both *FASN* and *ACACA* (Figure 2f). *C12orf49* also showed negative GIs with *LDLR*, *ACSL1* (i.e. encoding acyl-CoA synthase), *SLC25A1* (i.e. encoding mitochondrial citrate transporter), *SCD* and *SREBF2* further supporting a role for this gene in fatty acid biosynthesis (Figure 2f). Consistently, *C12orf49* negative GIs were enriched for genes involved in fatty acid metabolism, cholesterol biosynthesis and additional metabolic pathways (FDR <0.05) (Figure 4a, **Table S3).** Moreover, as observed for the *FASN* GI profile, *C12orf4*9 negative GIs involved genes functioning in vesicle-mediated trafficking and endocytosis, including *RAB3GAP2, RABIF, RAB18, VPS18, VPS419* and *VPS39* (**Table S2)**. Beyond vesicle trafficking, many of the genes that showed a negative GI with *C12orf49* also displayed negative GIs with other query genes in our lipid metabolism panel (e.g. *LDLR*, *ALG3, ASCL1, MBTPS2, SLC25A1, PDHA1*), supporting the functional relatedness of these genes (Figures 4b-c, S4f-h). Thus, our lipid metabolism GI map strongly implicates *C12orf49* as playing a functional role in lipid metabolism.

**Figure 4.**
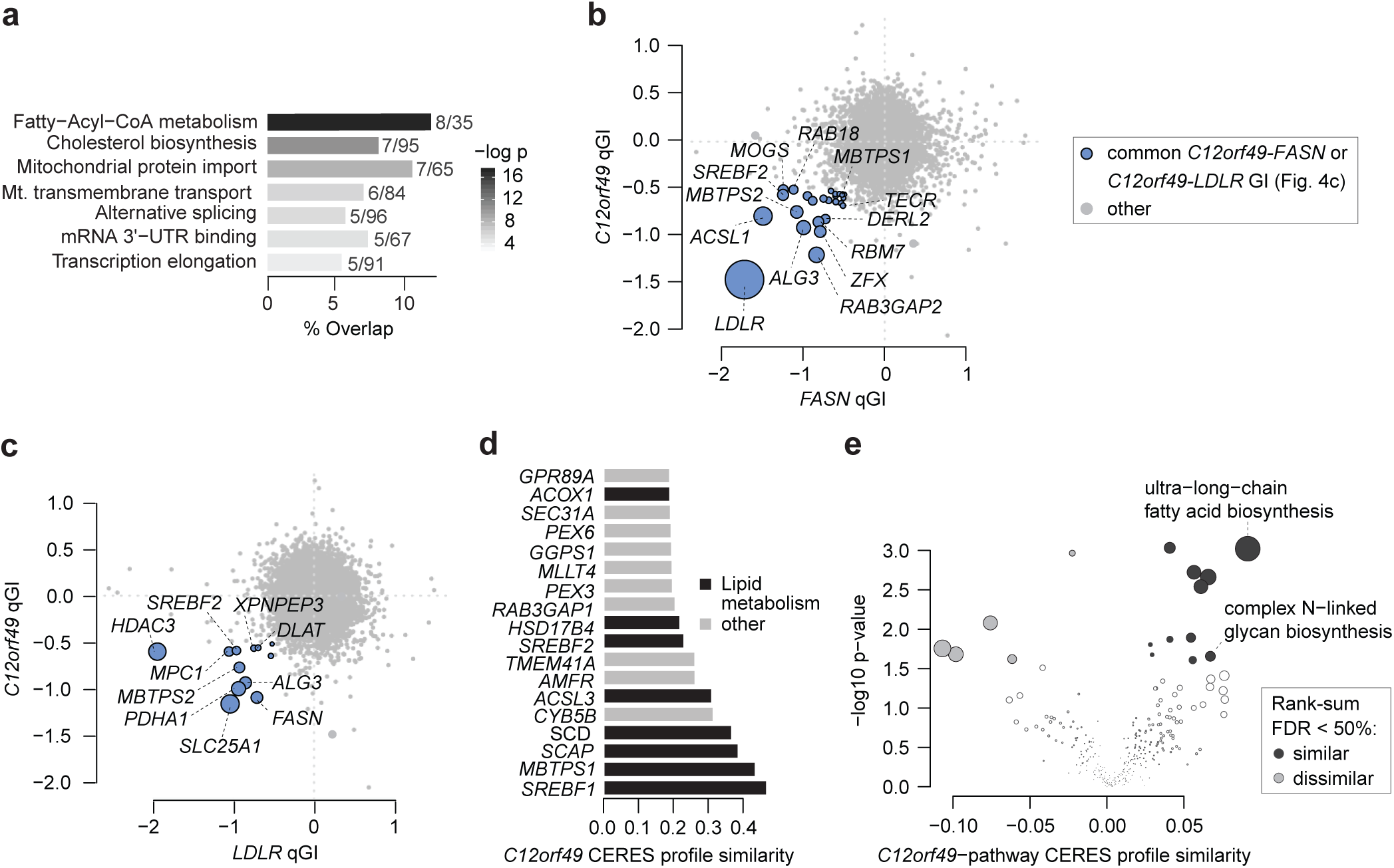
*C12orf49* genetic interaction profile suggests a functional role in lipid metabolism. (**a**) Bar plot depicting pathway enrichment of negative genetic interactions with *C12orf49* (|qGI| > 0.5, FDR < 0.5) using GO molecular functions, GO bioprocesses and Reactome standards. Significantly enriched gene sets (p < 0.05, maximum term size 100). Bars depict mean percentage overlap with the indicated term, and the numbers on each bar indicate the number of genes overlapping a particular term and term size, respectively. The greyscale color legend for p-values is indicated on the right. (**b**) Scatter plot of *C12orf49* and *FASN* qGIs depicting GI overlap between *C12orf49* and *FASN* qGI scores. *FASN* qGI scores are represented as the mean between three independent screens. A common negative GI is called if it is significant (qGI < −0.5, FDR < 0.5) in the *C12orf49*-KO screen and significant in 2 of 3 *FASN*-KO screens (indicated in blue). The top 10 strongest common GIs, lipid metabolism and vesicle trafficking genes are labelled. (**c**) Scatter plot of *C12orf49* and *LDLR* qGIs depicting GI overlap between *C12orf49* and *LDLR* qGI scores. A common negative GI is called if it is significant (qGI < −0.5, FDR < 0.5) in both screens (indicated in blue). The top 10 strongest common GIs and lipid metabolism genes are labelled. (**d**) Bar plot indicating the *C12orf49* profile similarity across genome-wide DepMap CRISPR/Cas9 screens. Similarity (i.e. co-essentiality) was quantified by taking all pairwise gene-gene Pearson correlation coefficients of CERES score profiles across 563 screens (19Q2 DepMap data release). The top 18 out of 17,633 gene profiles most similar to *C12orf49* are shown. Genes associated with lipid metabolism are indicated in black. (**e**) Volcano plot of pathway enrichment for *C12orf49* co-essential genes. *C12orf49* co-essentiality profile scores for all 17,634 genes represented in the DepMap were mean-summarized by pathway as defined in the HumanCyc standard (Romero *et al.*, 2004). Tendencies towards pathway-level similarity (co-essentiality) and dissimilarity (exclusivity) with C12orf49 were tested using a two-sided Wilcoxon rank-sum test followed by multiple hypothesis correction with the Benjamini and Hochberg procedure.

To further confirm the predictions about *C12orf49*’s function based on our HAP1 GI data, we also examined publicly available data from the 19Q2 DepMap release and observed that *C12orf49* is essential for fitness in 120 out of 563 cell lines with highest dependencies observed for lung, ovarian, pancreatic, colon and bile duct origins (Meyers *et al*, 2017; Behan *et al*, 2019). Other genes that shared similar cell line essentiality profiles to *C12orf49* included *SREBF1*, *SREBF2*, *MBTPS1*, *SCAP*, *SCD* and *ACSL3* (Figures 4d, S4i). The association of *C12orf49* with lipid metabolism genes was corroborated by a pathway enrichment analysis of the co-essentiality profiles, which revealed strong enrichment for genes annotated to ultra-long-chain fatty acid biosynthesis (Figures 4e, S4j). Interestingly, germline variants in *C12orf49* have also been reported to associate with serum lipid abnormalities in high-density lipoprotein (HDL) in a multi-ethnic cohort of the Million Veteran Program, further supporting a role for this gene in lipid metabolism (Klarin *et al*, 2018). Overall, these observations support a novel function for *C12orf49* in lipid metabolism that is conserved across diverse cell types.

### C12orf49 is a novel regulator of lipid uptake

We performed proximity-based labelling of proteins coupled to mass spectrometry (BioID-MS) to reveal potential C12orf49 protein interactions. Because the C12orf49 single predicted N-terminal transmembrane domain may direct the C-terminal DUF2054 domain into the lumen of the secretory pathway, leaving the N-terminus facing the cytoplasm, BioID-MS was performed separately with both N- and C-terminal BirA-tagged C12orf49 open reading frames (ORFs) expressed in HEK293 cells. Proximity-based labelling with the N-terminal construct captured proteins localizing to various cellular compartments including the ER, Golgi apparatus, plasma membrane and the cytosol, whereas the C-terminal BirA construct revealed a strong enrichment of proteins localizing to the endoplasmic reticulum (ER) lumen (Figure 5a, **Table S5**). Furthermore, the BirA ligase fused to the N-terminal BirA ligase captured proximal interactions with proteins functioning in vesicle and ER – Golgi transport, whereas C-terminus labelled proteins enriched for functions related to protein folding and glycosylation (Figure 5b, **Table S6**). Together, these results further support that C12orf49 localizes to the ER membrane or transport vesicles that may traffic to or from the ER, whereby its N-terminus likely faces the cytoplasm and, in this context, the C-terminus would face the ER lumen.

**Figure 5.**
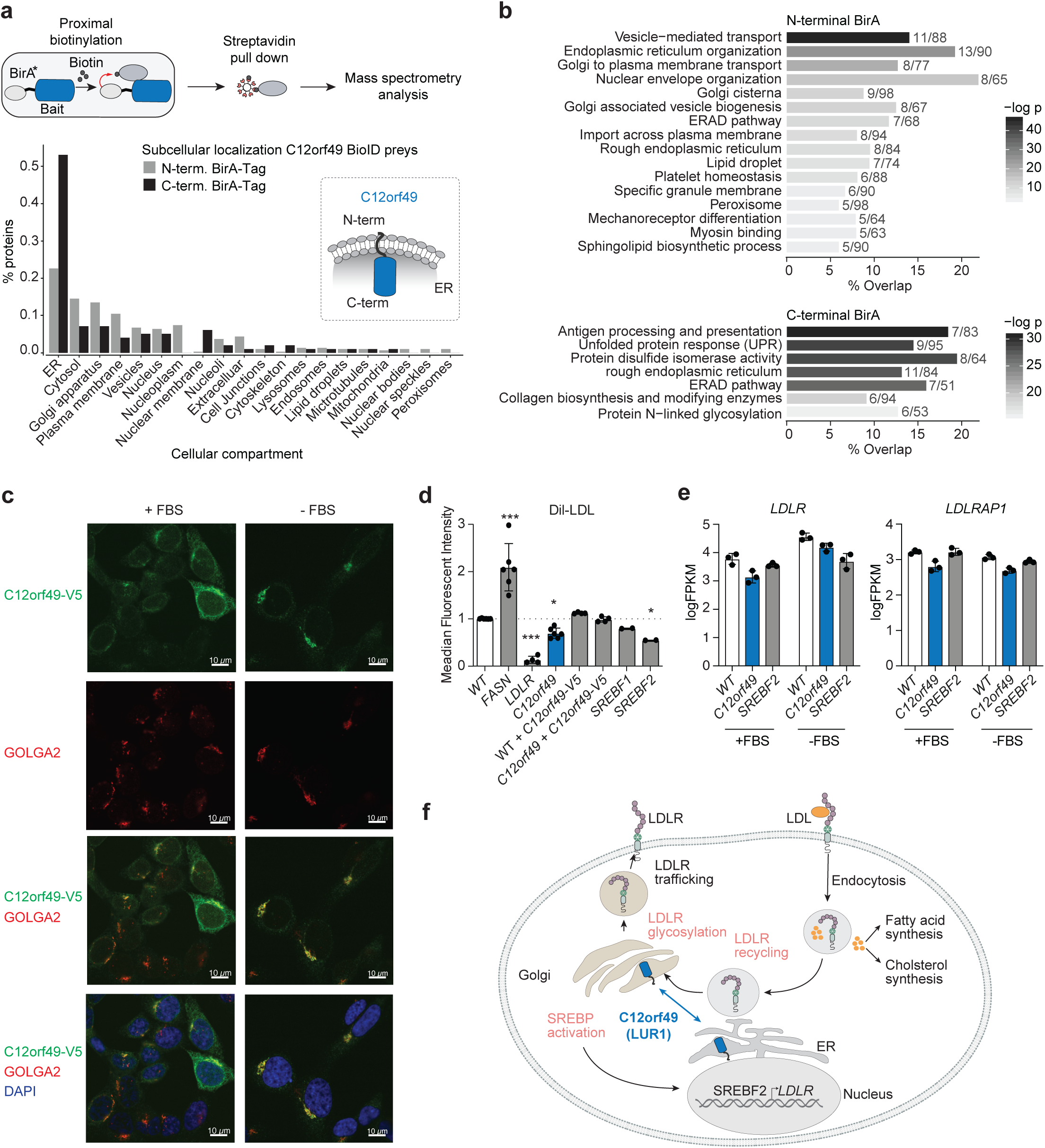
C12orf49 shuttles between ER and Golgi and regulates lipid uptake. (**a**) Schematic outlining proximal protein capture using BioID mass spectrometry analysis (upper panel) and analysis of subcellular localization of C12orf49 BioID preys (lower panel). Barplots depicting the fraction of proteins localizing to indicated cellular compartments for preys captured with N-terminal (grey) or C-terminal (black) BirA*-tagged C12orf49 in 293 cells. The inset shows a schematic representation of the predicted topology and orientation of C12orf49 with respect to the cytoplasm and ER. (**b**) Pathway enrichment analysis of BioID preys captured with N-terminal (top panel) or C-terminal (bottom panel) BirA*-tagged C12orf49 using the GO molecular function, biological process and Reactome standards. Terms for significantly enriched gene sets (p < 0.05, maximum term size 100) are indicated and bars depict mean percentage overlap with the indicated term. Numbers indicate the number of genes overlapping a particular term and term size, respectively. The greyscale color legend for p-values is indicated on the right. (**c**) Immunofluorescence microscopy analysis of C-terminal V5-tagged C12orf49 in HAP1 cells under normal (left) or serum-starved (right) growth condition. C12orf49-V5 localization is shown in green, GOLGA2 is a marker of the Golgi apparatus and shown in red, and DAPI (blue) marks the nuclei. Scale bars correspond to 10 µm. (**d**) Bar plots showing the results of low density lipoprotein (LDL) uptake assays in the indicated cells using the Dil-LDL probe. All data are represented as means ± standard deviation (n = 2-6). ***p < 0.001 and *p < 0.05; one-way ANOVA. (**e**) Bar plots indicating FPKM expression values from RNA sequencing data for *LDLR* and *LDLRAP1* in WT, *C12orf49*-KO, and *SREBF2*-KO cells under normal (+FBS) and serum-starved (-FBS) growth conditions as assessed by RNA sequencing (n=3). (**f**) Model summarizing functions and locations of key players in lipid metabolism, including LUR1 (C12orf49).

We performed immunofluorescence analysis to study the subcellular localization of C12orf49 under normal and starved conditions. Under normal growth conditions (with serum), C12orf49 containing a C-terminal V5 tag (i.e. C12orf49-V5) was localized throughout the ER-Golgi network (Figure 5c), consistent with our BioID results. Strikingly, C12orf49-V5 accumulated in the Golgi apparatus under serum starvation, as assessed by co-staining with GOLGA2, a Golgi membrane marker protein (Figure 5c). These data thus suggest that localization of C12orf49 is regulated in a growth condition-dependent manner, involving the shuttling between the ER and the Golgi apparatus.

Together, genetic and proteomic interaction data indicate that C12orf49 may play a role in lipid metabolism and vesicle-mediated transport. To explore this hypothesis, we measured uptake of labelled LDL particles, which represent one of the major sources of extracellular fatty acids, across several HAP1 KO lines. As expected, loss of *LDLR* resulted in abolishment of LDL-staining, while *FASN*-KO cells displayed increased uptake of exogenous lipid (Figures 5d, S5a). In contrast, loss of *C12orf49* caused a significant reduction of LDL uptake, which was rescued by the exogenous expression of C12orf49 (Figures 5d, S5a). Because since *C12orf49*-KO cells do not exhibit reduced uptake of labelled transferrin (Figure S5b), the reduction of LDL uptake is specific to lipid transport and is not a consequence of a general defect in receptor-mediated endocytosis,. Interestingly, a similar reduction in LDL uptake was also observed in *SREBF1*- and *SREBF2*-deficient cells. Overall, these results indicate that *C12orf49* impacts LDL uptake and support the GIs identified between *C12orf49* and genes functioning in fatty acid biosynthesis and lipid homeostasis.

Sterol regulatory element-binding proteins (SREBPs) traffic to the Golgi where they are cleaved such that the processed form enters the nuclease to activate transcription of genes regulating lipid homeostasis (Brown & Goldstein, 1997; Horton *et al*, 2008). Thus, C12orf49 could somehow play a role in the activation of the SREBP transcription factors. To explore this possibility, we performed RNA-sequencing experiments under normal and serum-starved conditions across HAP1 WT, *C12orf49*-KO and *SREBF2*-KO cells (**Table S7**). As expected, serum-starvation resulted in induction of a cholesterol biosynthetic transcriptomic signature in HAP1 WT cells but not in *SREBF2*-KO cells (Figures 5e, S5c-d). In *C12orf49*-KO cells, we observed a SREBP-mediated transcriptional response similar to WT cells, suggesting that C12orf49 is not absolutely required for the activation of SREBP upon serum starvation (Figure 5e, S5c-d). However, we did notice a trend for lower expression of cholesterol biosynthesis and LDL uptake genes in *C12orf49*-KO cells, which was confirmed by qRT-PCR (Figure 5e, S5e). The absence of strong gene expression changes suggests that C12orf49 may also regulate LDL uptake on a post-transcriptional level. We therefore measured LDLR protein levels on the cell surface by flow cytometry but we did not observe any significant changes in LDLR localization or abundance in *C12orf49*-KO compared to WT cells (Figure S5f). Thus, C12orf49 may influence LDL uptake through the regulation of post-translational modifications of LDLR, such as glycosylation, which takes place in the ER and Golgi apparatus (Figure 5f).

In summary, our unbiased GI screens and follow-up experiments have revealed that the uncharacterized gene *C12orf49* plays a role in the regulation of lipid transport and our data further indicate that its subcellular localization is dynamically regulated in a growth condition-dependent manner throughout the ER-Golgi network. Our findings indicate that C12orf49 mainly regulates lipid uptake on a post-transcriptional level and we suggest that *C12orf49* be named *LUR1* for its role in Lipid Uptake Biology. We speculate that the *LUR1* product may be involved in some aspect of the glycosylation of LDLR, the recycling of vesicles to the cell surface, or in regulating the transcriptional response mediated by SREBPs (Figure 5f).

## DISCUSSION

The systematic mapping of GIs in model organisms like yeast has provided a detailed view into the functional organisation of eukaryotic cells (Costanzo *et al*, 2019). Recent advances in CRISPR-based genome engineering technologies provide a path for similar systematic GI studies in human cells (Horlbeck *et al*, 2018; Najm *et al*, 2018; Han *et al*, 2017; Norman *et al*, 2019; Shen *et al*, 2017). Here, we apply genome-wide CRISPR-based fitness screens using query mutant HAP1 cell lines to systematically map GIs with a focus on lipid metabolism. Our data revealed a strong interaction between *de novo* fatty acid synthesis and lipid uptake processes, highlighting a system that balances synthesizing lipids intracellularly with their uptake from the extracellular environment. More generally, this analysis confirms that relatively strong negative GIs identify functionally related genes, mapping a functional wiring diagram for a particular cellular process.

We screened a *FASN* mutant query cell line multiple times and identified highly confident negative GIs, many of which were involved in lipid metabolism. Perturbation of *de novo* fatty acid synthesis has been suggested as a prominent cancer therapeutic approach and multiple compounds targeting FASN are currently being tested in clinical trials; for example, TVB-2640 is a FASN inhibitor that is being tested in solid tumors in phase 2 trials, while both Fatostatin and Betulin are inhibitors of the SREBP-SCAP interaction in pre-clinical development (Röhrig & Schulze, 2016; Brenner *et al*, 2017). Since single agent therapies often lead to emergence of resistance and tumor relapse, it makes sense to pursue therapeutic targets that are synergistic with FASN inhibition. Thus, the strong GIs detected in our FASN screen may be informative towards future investigations of combinatorial targets or biomarkers to treat diseases that would benefit from disruption of *de novo* fatty acid biosynthesis.

Our focused GI landscape related to *de novo* fatty acid biosynthesis provides unique insight into the genetic dependencies required for response to perturbation of lipid metabolism. Several pathways emerge as being most commonly utilized to adapt to perturbations, including those involved in alternate fatty acid and cholesterol biosynthesis processes as well as lipid uptake. Interestingly, while our screens revealed strong negative GIs between *de novo* fatty acid synthesis and uptake of LDL, we failed to detect interactions with transporters of fatty acids. This may be a consequence of the genetic redundancy inherent amongst the *SLC27A* (FATP) fatty acid transporter family (Gimeno, 2007). As previously shown in yeast (VanderSluis *et al*, 2010), functional redundancy between paralogs can mask genetic interactions associated with perturbation of a single gene of a duplicated pair and highlights an important need for multi-gene targeting systems to survey complex genetic interactions involving more than two genes. Nonetheless, our data suggest a strong functional relationship between *de novo* fatty acid synthesis and glycosylation, and may involve a mechanism wherein cells modify the FATP transporters through N-glycosylation, thereby enhancing lipid uptake as suggested by Ryczko *et al* (Ryczko *et al*, 2016). As such, this pathway serves as an obvious focal point not only for ongoing mechanistic investigation but also therapeutic development for anti-cancer strategies targeting *de novo* fatty acid synthesis.

Genome-wide GI profiling also revealed an important role for *LUR1* (*C12orf49)* in lipid uptake. Interestingly, analysis of the DepMap data revealed that *LUR1* is essential in the same set of cancer cell lines that also depend on other lipid biosynthesis-related genes for viability, including *SREBF1*, *MBTPBS1*, *SCAP* and *SCD*. Similarly, two recent studies identifying co-functional gene clusters, support a functional role of *LUR1* in lipid metabolism across diverse genetic backgrounds (Boyle *et al*, 2018; Kim *et al*, 2019). Furthermore, genome-wide association studies with large patient cohorts have found *LUR1* variants linked to abnormal HDL profiles (Klarin *et al*, 2018), neuroticism (Luciano *et al*, 2018; Kichaev *et al*, 2019; Nagel *et al*, 2018), body height (Kichaev *et al*, 2019), and neuroticism (Nagel *et al*, 2018), all phenotypes that could have root causes in lipid metabolism defects.

In summary, we provide an unbiased and genome-wide approach for uncovering genetic vulnerabilities related to lipid metabolism in human cells, which led us to identify a function for *LUR1*. Our GI profiles for *de novo* fatty acid synthesis and related lipid uptake genes provide a resource for studying metabolic rewiring and disease phenotypes linked to lipid metabolism. We also demonstrate the power of systematic GI profiling using query mutants in a coisogenic cell line, an approach that can be applied to other bioprocesses and expanded to begin generating more comprehensive GI maps for human genes.

## ACKNOWLEDGMENTS

We thank members of the Moffat lab for helpful discussions. Queenie Huang and Shaan Sidhu are gratefully acknowledged for assistance with molecular biology experiments. M.A. was supported by a Swiss National Science Foundation Postdoctoral Fellowship; K.A.L. was supported by a Vanier Canada Graduate Scholarship and Studentship award from the Kidney Cancer Research Network of Canada. M.B. was supported by a DFG Fellowship (Bi 2086/1-1). This research was funded by grants from the Canadian Institutes for Health Research (J.M., C.B and B.A), Ontario Research Fund (B.A., C.B. and J.M) and Canada Research Chairs Program (J.M. and C.B.). C.L.M., M.B., and M.R. are partially supported by grants from the National Science Foundation (MCB 1818293) and the National Institutes of Health (R01HG005084, R01HG005853).

## AUTHOR CONTRIBUTIONS

Conceptualization and design of the study: M.A., K.A.L., and J.M.; Experimental investigation: M.A., K.A.L., A.H.Y.T., K.C., L.N., O.S., A.H., J.P., Z.L., H.A. and A.W. Data analysis: M.A., K.L., M.B., M.C., M.R., K.R.B., C.R., M.U., P.M., J.W.D, A.C.G, J.L.M and J.M.; Writing & Editing: M.A., K.A.L., M.B., M.C., B.J.A., C.B., C.L.M. and J.M. with the input from other authors; Supervision: J.W.D., A.C.G., C.L.M., B.J.A., C.B., and J.M.; Funding Acquisition: C.L.M., B.J.A., C.B. and J.M..

## METHODS

### Cell culture

Human HAP1 wild type cells were obtained from Horizon Genomics (clone C631, sex: male with lost Y chromosome, RRID: CVCL_Y019). The following HAP1 gene knockout cell lines were obtained from Horizon: *FASN* (HZGHC003700c006), *ACACA* (HZGHC004903c002), *LDLR* (HZGHC003978c007), *SREBF1* (HZGHC001361c012), *SREBF2* (HZGHC000683c004). All gene knockout cell lines were confirmed to carry the expected out-of-frame insertions or deletions by Sanger Sequencing of PCR products. HAP1 cells were maintained in low glucose (10 mM), low glutamine (1 mM) DMEM (Wisent, 319-162-CL) supplemented with 10% FBS (Life Technologies) and 1% Penicillin/Streptomycin (Life Technologies). This culture medium is referred to as “minimal medium”. Cells were dissociated using Trypsin (Life Technologies) and all cells were maintained at 37°C and 5% CO_2_. Cells were regularly monitored for mycoplasma infection.

### HAP1 KO cell line generation

The HAP1 *C12orf49* gene knockout cell line was constructed by first cloning a gRNA targeting C12orf49 (Table S8) into the pX459v2 backbone (Addgene #62988), which was modified to carry the same restriction overhangs as the pLCKO vector (Addgene #73311). 350k HAP1 WT cells were seeded into a 6-well plate and 24 hours later cells were transfected with a mix of 2 µg pX459 plasmid (Addgene #62988) carrying a gRNA, 6 µl X-treme Gene transfection reagent (Roche), and 100 µl Opti-MEM media (Life Technologies). Twenty-four hours after transfection, cells were selected in medium containing 1 µg/ml puromycin for three days and single cells were sorted onto 96-well plates by manual seeding of a single cell suspension at 0.6 cells/well. Following amplification of cells from individual wells, genomic DNA was extracted with Extracta DNA Prep (Quanta Bio), Sanger sequencing was performed across the gRNA target sites following PCR amplification, and successful gene knockouts were identified following sequence analysis.

### Library virus production and MOI determination

For CRISPR library virus production, 8 million HEK293T cells were seeded per 15 cm plate in DMEM medium containing high glucose, pyruvate and 10% FBS. Twenty-four hours after seeding, the cells were transfected with a mix of 8 µg lentiviral lentiCRISPRv2 vector containing the TKOv3 gRNA library (Addgene #90294) (Hart *et al*, 2017), 4.8 µg packaging vector psPAX2, 3.2 µg envelope vector pMD2.G, 48 µl X-treme Gene transfection reagent (Roche) and 1.4 ml Opti-MEM media (Life Technologies). Twenty-four hours after transfection, the media was replaced with serum-free, high-BSA growth media (DMEM, 1.1g/100ml BSA, 1% Penicillin/Streptomycin). Virus-containing media was harvested 48 hours after transfection, centrifuged at 1,500 rpm for 5 minutes, aliquoted and frozen at −80°C.

For determination of viral titers, 3 million HAP1 cells seeded in 15 cm plates were transduced with different dilutions of the TKOv3 lentiviral gRNA library along with polybrene (8 µg/ml), in a total of 20 ml medium. After 24 hours, the virus-containing media was replaced with 25 ml of fresh media containing puromycin (1 µg/ml), and cells were incubated for an additional 48 hours. Multiplicity of infection (MOI) of the titrated virus was determined 72 hours post-infection by comparing percent survival of puromycin-selected cells to cells that were infected but not selected with puromycin (i.e. puro minus controls).

### Pooled CRISPR dropout screens

For pooled CRISPR dropout screens, 3 million HAP1 cells were seeded in 15 cm plates in 20 ml of specified media. A total of 90 million cells were transduced with the lentiviral TKOv3 library at a MOI∼0.3, such that each gRNA is represented in about 200-300 cells. Twenty-four hours after infection, transduced cells were selected with 25 ml medium containing 1 µg/ml puromycin for 48 hours. Cells were then harvested and pooled, and 30 million cells were collected for subsequent gDNA extraction and determination of the library representation at day 0 (i.e. T0 reference). The pooled cells were then seeded into three replicate plates, each containing 18 million cells (>200-fold library coverage), which were passaged every three days and maintained at >200-fold library coverage until T18. Genomic DNA pellets from each replicate were collected at each day of cell passage.

### Preparation of sequencing libraries and Illumina sequencing

Genomic DNA was extracted using the Wizard Genomic DNA Purification Kit (Promega). The gDNA pellets were resuspended in TE buffer, and the concentration was estimated by Qubit using dsDNA Broad Range Assay reagents (Invitrogen). Sequencing libraries were prepared from 50 µg of the extracted gDNA in two PCR steps, the first to enrich guide-RNA regions from the genome, and the second to amplify guide-RNA and attach Illumina TruSeq adapters with i5 and i7 indices as described previously using staggered primers aligning in both orientations to the guide-RNA region (Table S8) (Aregger *et al*, 2019). Barcoded libraries were gel purified and final concentrations were estimated by quantitative RT-PCR. Sequencing libraries were sequenced on an Illumina HiSeq2500 using single read sequencing and completed with standard primers for dual indexing with HiSeq SBS Kit v4 reagents. The first 21 cycles of sequencing were dark cycles, or base additions without imaging. The actual 36-bases read begins after the dark cycles and contains two index reads, reading the i7 first, followed by i5 sequences. The T0 and T18 time point samples were sequenced at 400- and 200-fold library coverage, respectively.

### Construction of color-coded lentiCRISPRv2 vectors for co-culture assay

The color-coded lentiCRISPRv2 vectors were derived from the lentiCRISPRv2 vector (Addgene #52961) by inserting mCherry (Addgene #36084) or mClover3 (Addgene #74236) open reading frames between the Cas9 and PuroR expression cassette. To this end, the lentiCRISPRv2 vector was digested with BamHI, PCR products coding for the respective fluorescent protein flanked by T2A and P2A self-cleaving peptides were ligated into the vector using Gibson assembly. The two forward primers (Table S8) were used at a 1:0.1:1 (P233:P234:P235) ratio in the same PCR reaction with the reverse primer (primers bind to both fluorescent proteins mCherry and mClover3).

### Validation of genetic interactions using co-culture assays

For validation of genetic interactions, HAP1 parental and gene knockout clones were transduced with color-coded lentiCRISPRv2 vectors targeting either an intergenic site in the AAVS1 locus (i.e. negative control), or a specific target gene hit (e.g. *LDLR*). Each gene was targeted with three independent and unique gRNAs. Twenty-four hours after transduction, cells were selected with 1 µg/ml puromycin for 48 hours and seeded for co-culture proliferation assays as follow: 50k of green (e.g. lentiCRISPRv2-mClover3 *AAVS1* gRNA) and red (e.g. lentiCRISPRv2-mCherry hit gene gRNA) cells were mixed (total 100k) in a 6-well plate in both color orientations for both parental and gene knockout cells, respectively. Cells were passaged every 4 days until day 12 (T12). Cells were trypsinized, washed and stained for dead cells using Zombie NIR (BioLegend). The relative proportion of red and green cells in the co-culture were assessed using an LSR Fortessa flow cytometer (BD Bioscience). The relative ratio of Hit:AAVS1 was calculated and averaged for the three gene-targeting guides and two color orientations.

### Low-density lipoprotein and transferrin uptake assay

For uptake experiments with labelled probes 150k HAP1 cells were seeded in a 12-well plate. After 48 hours cells were serum-starved overnight in minimal medium (described above) complemented with 0.3% BSA (BioShop) instead of FBS. After 16 hours cells were labelled with Dil-LDL (Invitrogen L3482), pHrodo Red LDL (Invitrogen L34356) or pHrodo Red Transferin (P35376) at 2 µg/ml (1:500) in minimal medium plus 0.3% BSA for 15 minutes at 37°C. Cells were washed in PBS, trypsinized and stained with 7-AAD (BioLegend 420404) or Zoombie NIR (BioLegend 423105) cell viability solution at 25 ng/ml (1:2,000) for 5 minutes at room temperature. Staining was measured using an LSR Fortessa flow cytometer (BD Bioscience).

### Proximity-based labelling of proteins capture to mass spectrometry (BioID-MS)

BioID-MS analysis was performed essentially as described previously (Hesketh *et al*, 2017), with minor modifications. In brief, HEK293 Flp-In T-REx lines expressing inducible N- or C-terminal BirA*-FLAG-tagged C12ORF49 open reading frames were generated. Cells were treated with 1 µg/ml tetracycline to induce expression of baits and 50 µM biotin for labelling of proximal proteins. After 24 hours cell pellets were collected and lysed in RIPA lysis buffer (50mM Tris-HCl pH 7.5, 150mM NaCl, 0.1% (w/v) SDS, 1% NP-40, 1mM EDTA, 1mM MgCl_2_; 0.5% Deoxycholate and Sigma protease inhibitors were added right before cell lysis.) at an 1:10 (g:ml) ratio, sonicated three times for 5 seconds with 2 seconds breaks. 1ul/sample TurboNuclease (BioVision) and 1ul/sample RNAse (Sigma) was added and samples were incubated at 4°C for 30 minutes. 20% SDS was added to bring the sample’s final SDS concentration to 0.25%, samples were mixed well and centrifuged at 14,000 rpm (Microfuge) for 20 mins in 4°C. The supernatant was added to Streptavidin resin (pre washed with lysis buffer) using 30µl bed volume and rotated at 4°C for 3 hours. Beads were washed after binding as following: a) 1×1ml of 2% SDS buffer (2% SDS, 50mM Tris-Hcl pH7.5), b) 1×1ml of lysis buffer, c) 1×1ml of HEK293 lysis buffer (with 0.1% NP-40), d) 3×1ml of 50mM ammonium bicarbonate (made fresh). After purification of biotinylated preys using streptavidin sepharose, samples were digested on beads using trypsin. Samples were separated by liquid chromatography and analysed by tandem mass spectrometry on a Thermo Orbitrap Elite mass spectrometer. Data processing and analysis was performed within the ProHits LIMS (Liu *et al*, 2016) searched against the RefSeq human and adenovirus data base, version 57; forward and reverse. Mascot and Comet search results were jointly analysed using the iProphet component of the Trans Proteomic Pipeline. High confidence interactions were determined by scoring bait samples against negative control samples (8 negative controls consisting of either BirA*-FLAG alone, BirA*-FLAG-EGFP, empty vector backbone or EGFP alone were analysed; twelve samples for different baits, SLCO4A1, SLC35A1, UAP1L1 and C1orf115, were also included in this analysis) using the statistical tool SAINTexpress v3.6.1 with two-fold compression of the negative controls and default parameters). Preys with a SAINT score (FDR) of less than 1% were considered as high confidence hits. All mass spectrometry data will be deposited to ProteomeXchange through partner MassIVE (massive.ucsd.edu) upon publication of manuscript.

### Western Blotting

HAP1 WT and *FASN* KO cells were lysed in buffer F (10 mM Tris pH 7.05, 50 mM NaCl, 30 mM Na pyrophosphate, 50 mM NaF, 10% Glycerol, 0.5% Triton X-100) and centrifuged at 14,000 rpm for 10 minutes. The supernatant was collected and protein concentration was determined using Bradford reagent (BioRad). 10 µg protein was resolved on 4 - 12% Bis-Tris gels (Life Technologies) and transferred to Immobilon-P nitrocellulose membrane (Millipore) at 66V for 90 minutes. Subsequently, proteins were detected using anti-FASN (1:2,000, Abcam ab128870) and anti-β-Actin (1:10,000, Abcam ab8226) antibodies and proteins were visualized on X-ray film using Super Signal chemiluminescence reagent (Thermo Scientific).

### Immunofluorescence

Cells were seeded on cover slips and fixed with 4% paraformaldehyde in PBS for 10 minutes at room temperature. Cells were permeabilized with 1% NP-40 in antibody dilution solution (PBS, 0.2% BSA, 0.02% sodium azide) for 10 minutes and blocked with 1% goat serum for 45 minutes. Cells were incubated with anti-V5 (1:250, Abcam ab27671) and anti-GOLGA2 antibodies (1:250, Sigma HPA021799) for 1 hour at room temperature. Subsequently, cells were incubated with Alexa Fluor488 goat anti-mouse (1:500, Invitrogen A-11001) or Alexa Fluor647 anti-rabbit antibodies (1:500, Invitrogen A-21245) and counterstained with 1 µg/ml DAPI (Cell Signaling Technology, 4083S) for 45 minutes in the dark. Cells were visualized by confocal microscopy (Zeiss LSM 880).

### Protein expression analysis by flow cytometry

Cells were detached using accutase (GIBCO), washed in PBS and 250k cells were stained with PE-LDLR at 2 µg/ml (1:100, BD Bioscience 565653) for 20 minutes at 4°C and Zoombie NIR (BioLegend 423105) cell viability solution at 25 ng/ml (1:2,000) for 5 minutes at room temperature. Staining was measured using an LSR Fortessa flow cytometer (BD Bioscience).

### RNA-sequencing

#### Sample preparation

HAP1 WT, *FASN* KO and *C12orf49* KO cells were cultured in minimal DMEM medium for 48h and either control treated or serum-starved for 4 hours as indicated. Each cell line was cultured and processed in three biological replicates. RNA was extracted using the RNeasy Kit (QIAGEN) according to manufacturer’s instructions. 18 total RNA samples were DNase treated using RNase-free DNase Set (Qiagen, 79254). Samples were submitted for mRNA-Seq at the Donnelly Sequencing Centre at the University of Toronto (http://ccbr.utoronto.ca/donnelly-sequencing-centre). RNA was quantified using Qubit RNA BR (Thermo Fisher Scientific, Q10211) fluorescent chemistry and 1 ng was used to obtain RNA Integrity Number (RIN) using the Bioanalyzer RNA 6000 Pico kit (Agilent Technologies, 5067-1513). Lowest RIN was 9.5; median RIN score was 9.8. 1000 ng per sample was then processed using the NEBNext Ultra II Directional RNA Library Prep Kit for Illumina (New England Biolabs, E7760L) and included polyA-enrichment using NEBNext Poly(A) mRNA Magnetic Isolation Module (New England Biolabs, E7490L), fragmentation for 15 minutes at 94°C prior to first strand synthesis, and 8 cycles of amplification after adapter ligation. 1µL top stock of each purified final library was run on an Agilent Bioanalyzer dsDNA High Sensitivity chip (Agilent Technologies, 5067-4626). The libraries were quantified using the Quant-iT dsDNA high-sensitivity (Thermo Fisher Scientific, Q33120) and were pooled at equimolar ratios after size-adjustment. The final pool was run on an Agilent Bioanalyzer dsDNA High Sensitivity chip and quantified using NEBNext Library Quant Kit for Illumina (New England Biolabs, E7630L). The quantified pool was hybridized at a final concentration of 400 pM and sequenced paired-end on the Illumina NovaSeq6000 platform using a S2 flowcell at 2×151 bp read lengths.

#### Data Processing

Samples were mixed to obtain an average of 35 million clusters that passed filtering. Reads shorter than 36bp on either read1 or read2 were removed prior to mapping. Reads were aligned to reference genome hg38 and Gencode V25 gene models using the STAR short-read aligner (v2.6.0a) (REF). Approximately 80% of the filtered reads mapped uniquely, and the read counts from each sample, computed by STAR, were merged into a single matrix using R. The raw and processed data will be deposited in the GEO database upon publication of manuscript.

#### Differential expression

Differentially expressed genes were identified using the Bioconductor packages limma (v3.32.10) and edgeR (v3.24.3). The read count matrix was filtered using the filterByExpr() function using default parameters. Principal Components Analysis was performed to examine the main treatment effects, and to exclude the presence of confounding batch effects, using the base R function prcomp(). Samples were normalized using calcNormFactors(method=”TMM”) from edgeR and transformed to log2 using voom(). Next, a design matrix was specified to fit coefficients for the CRISPR knockouts, presence or absence of FBS, and an interaction term to examined differences in the FBS effect in the mutant backgrounds. Differentially expressed genes were extracted using topTable() with log2(fold-change) > 0.58 and adjusted P-value less than 0.05.

### Quantitative real-time (qRT)-PCR analysis

HAP1 WT, *FASN* KO and *C12orf49* KO cells were cultured in minimal DMEM medium for 48h and either control treated or serum-starved for 4 hours as indicated. RNA was extracted using the RNeasy Kit (QIAGEN) according to manufacturer’s instructions. RNA was converted into cDNA using the cVilo master mix (ThermoScientific) according to manufacturer’s instructions. The cDNA was amplified and quantified by quantitative PCR using the Maxima SYBR Green PCR master mix (ThermoScientific) according to manufacturer’s instructions. Transcript levels were normalized to *GAPDH* (see Table S8 for primer sequences).

### Metabolite profiling

HAP1 WT and *FASN*-KO cells were cultured in minimal medium for 3 days. Cells were washed twice in warm PBS and subsequently flash frozen on liquid nitrogen. Cells were scraped in chilled extraction solvent (40% Acetonitrile: 40% Methanol: 20% water, all HPLC grade), transferred to clean tubes and shaken for one hour at 4°C and subsequently centrifuged at 4°C at max speed for 10 minutes. The supernatants were transferred to a clean tube and dried in a speedvac then stored at −80°C until mass spec analysis. Samples were reconstituted in water containing Internal Standards D7-Glucose and _13_C_15_N-Tyrosine and injected twice through the HPLC (Dionex Corporation) for positive and negative mode analysis using a reverse phase column (Inertsil ODS-3, 4.6 mm internal diameter, 150 mm length, and 3 µM particle size). In positive mode analysis, the mobile phase gradient ramped from 5% to 90% acetonitrile in 16 minutes, remained for 1 minute at 90%, then returned to 5% acetonitrile in 0.1% acetic acid over two minutes. In negative mode, the acetonitrile composition ramped from 5 to 90% in 10 minutes, remained for 1 minute at 90%, then returned to 5% acetonitrile in mobile phase (0.1% tributylamine, 0.03% acetic acid, 10% methanol). The total runtime in both the positive and negative modes was 20 minutes, the samples were maintained at 4°C, and the injection volume was 10 µL. An automated washing procedure was included before and after each sample to avoid any sample carryover.

The eluted metabolites were analyzed at the optimum polarity in MRM mode on an electrospray ionization (ESI) triple-quadrupole mass spectrometer (ABSciex 5500 Qtrap). The mass spectrometric data acquisition time for each run was 20 minutes, and the dwell time for each MRM channel was 10 ms. Mass spectrometric parameters were as previously published (Abdel Rahman *et al.,* 2013). Metabolite peak areas were determined using Multiquant software (SCIEX, Toronto, ON, Canada), normalized to internal standard in each mode yielding an area ratio and then further normalized to total cell number for each sample and Malonyl-CoA levels were further normalized to WT cells.

## QUANTIFICATION AND STATISTICAL ANALYSIS

### Guide Mapping and Quantification

FASTQ files from single read sequencing runs were first trimmed by locating constant sequence anchors and extracting the 20 bp gRNA sequence preceding the anchor sequence. Pre-processed paired reads were aligned to a FASTA file containing the TKOv3 library sequences using Bowtie (v0.12.8) allowing up to 2 mismatches and 1 exact alignment (specific parameters: -v2 -m1 -p4 --sam-nohead). Successfully aligned reads were counted, and merged along with annotations into a matrix.

### Scoring of quantitative genetic interactions: the qGI score

To identify and quantify genetic interactions (GI), genome-wide CRISPR/Cas9 screens were performed using the TKOv3 gRNA library in HAP1 coisogenic cell lines. Coisogenic knockout (KO) “query” cell lines were obtained from Horizon Genomics (see above) or generated by introducing mutations in target genes of interest (see above) in the parental HAP1 cells, which we consider as wild-type (WT). The TKOv3 library contains 71,090 guide (g)RNAs that target ∼18k human protein-coding genes, most of them with four sequence-independent gRNAs (Hart *et al*, 2017). To quantify GIs, log_2_ fold-changes (LFC) between read-depth normalized gRNA abundance in the starting population (T0) and the endpoint (T18) were computed. Matched T0 measurement assured that differences between screens during library infection and Puromycin selection would not result in false positive GIs. Matched T0 were stabilized using the median across many T0 measurements (common T0), and those two estimates were combined in a weighted fashion to minimize correlation between GI scores and residual T0 (matched T0 – common T0). gRNA-level residual scores were derived for a given genetic background by estimating a non-interacting model between LFC values in this background and 21 WT HAP1 backgrounds. To do so, for each WT-KO screen pair the population of LFC values were M-A-transformed, which contrast the per-gRNA LFC difference M with per-gRNA mean A. A Loess regression was fitted, which was additionally locally stabilized by binning the data along A and considered equal bin sizes and equal numbers of data points in every bin. For each gRNA, this resulted in 21 residual scores, which represent the contrasts of a given KO with the 21 WT HAP1 screen. Under the assumption that genetic interactions are sparse and that experimental artifacts such as batch effects would introduce additional signal into the population of residual values, we computed a weighted mean of its 21 residual scores by giving a higher weight to WT HAP1 screens with lower absolute residual mean of all 71k gRNAs. We refer to the resulting value for each gRNA as the “guide-level” GI score. Those guide-level GI scores were further normalized. First, locally-defined shifts towards negative or positive scores were identified and normalized, based on genome location of the target genes. Next, to remove unwanted effects that would arise from screen-to-screen variability, we quantified guide-level GI scores for each of the 21 WT HAP1 screens by contrasting a given WT screen to the remaining WT screens (as described for the KO-WT comparison above). Patterns that explain substantial variance among these WT guide-level GI scores are likely to correspond with unwanted experimental artefacts. To remove these artefacts from the GI data, we performed singular value decomposition (SVD) on guide-level GI scores of the HAP1 WT screens only. We then projected guide-level GI scores onto the left singular vectors, and subtracted the resulting signal from the GI scores.

Finally, we computed gene-level genetic interaction scores. First, gRNAs were excluded when their guide-level GI profile disagreed with those of the remaining gRNAs against the same gene. Specifically, the mean within-gene guide-level GI profile Pearson correlation coefficient was computed. For the gRNA with the lowest value we tested if (i) the mean of all those four gRNA values for a given gene was above a selected threshold, which indicated that sufficient signal was present in the guide-level GI profiles, and (ii) the lowest value differed from this mean. All remaining guide-level GI values per gene were mean-summarized and their significance was computed using limma’s moderated t-test followed by Benjamini-Hochberg multiple testing correction.

### Screen reproducibility analysis

Reproducibility of the gRNA library screening data in FASN-KO cells was tested across three independent screens. The three screens were started from independent infections with lentivirus packaged gRNA library and performed as described above. To assess reproducibility of fitness effects, a log2 fold-change (LFC) quantifying the drop-out between T0 (after puromycin selection) and T18 (endpoint) was computed for each gene by mean-summarizing the respective four gRNA LFC values. The Pearson correlation coefficients (PCCs) were computed between LFC values of all three pairs of independent replicates.

Our experiments were designed to quantify fitness effect differences due to the introduction of a specific mutation into an otherwise isogenic background (i.e. GIs). To assess reproducibility of GIs, PCCs were computed between qGI values of all pairs of independent replicates.

To test reproducibility of genes, each gene’s contribution to the covariance between a pair of FASN-KO screens was computed and divided by the product of standard deviations of both given screens. The resulting three pairwise (for replicates A-B, A-C, B-C) gene-level scores were mean-summarized to a FASN qGI reproducibility score.

### Reproducibility analysis of FASN interactions

We used an MCMC-based approach to measure the reproducibility of *FASN* GIs. Specifically, we first independently scored the three independent *FASN* replicate screens and applied an FDR threshold of FDR 50% to generate positive and negative GI profiles for each of the three screens. MCMC was then used to jointly infer false negative and false positive rates, as well as a binary consensus *FASN* GI profile (separately for positive/negative GI). Then, using this consensus profile as a standard for evaluation (assuming pairs with posterior probability of interaction of > 0.5 as positives), we measured precision and recall statistics (averaged across the three screens) at two different cut-offs: a “standard” cut-off (absolute qGI score > 0.5 and FDR 50%) and a “stringent” cut-off (absolute qGI score > 0.7 and FDR 20%).

### Precision-recall analysis

To control quality of genome-wide gRNA screens, gene-level fitness effects were estimated by computing a log2 fold-change (LFC) quantifying the drop-out between T_0_ (after puromycin selection) and T_18_ (endpoint) for each gene and mean-summarizing the respective four gRNA LFC values. Gold-standard essential (reference) and non-essential (background) gene sets were taken from Hart et al., 2015(Hart *et al*, 2015) and Hart et al., 2017(Hart *et al*, 2017). For the identification of reference genes using LFC values of a given screen was assessed by computing precision over true positive statistics.

### Functional evaluation of genetic interactions

To calculate the enrichment of metabolic GIs in different functional standards, we separated the metabolic GIs in two different sets: all (background) GI scores and high confidence (reference) GI (FDR < 0.5, |qGI| >= 0.5). Then we calculated the fold enrichment of the reference set against the background set in a particular functional standard. First, we computed the overlap of metabolic GI pairs as co-annotations in the standard. Then we divided the overlap density of the background set into the overlap density of the reference set to determine the fold enrichment. Once we got the fold enrichments, we calculated p-values on the actual overlap counts of the reference and background sets according to hypergeometric tests. We used four different functional standards: Human functional network (Greene *et al*, 2015), GO biological processes (Ashburner *et al*, 2000), Pathway (Canonical pathways from (Liberzon *et al*, 2011)), and HumanCyc (Romero *et al*, 2005).

### Gene ontology enrichment analysis

Gene ontology (GO) enrichment analysis for the *FASN* and *C12orf49* GI screen and the BioID experiments were performed using the gProfileR R package using the GO-Bioprocesses, GO-Molecular Function and Reactome pathway standards. For the GI screens, enrichment analysis was performed for significant negative GIs (qGI < −0.5, FDR < 0.5), enriched pathways (p<0.05, maximum term size 100) with a similarity of > 50% were collapsed using the Cytoscape Enrichment Map function and the mean percentage overlap of hits within the term were visualized on a bar plot. For the BioID experiments, enrichment analysis was performed for significant hits (spectral counts > 10, FDR < 0.01), enriched pathways (p<0.05) with a similarity of > 50% were collapsed using the Cytoscape Enrichment Map function and the mean percentage overlap of hits within the term were visualized on a bar plot.

### Statistical Analysis

For all experiments the number of technical and/or biological replicates are listed in the figure legends or text. Unless otherwise indicated, statistical significance was assessed via one or two factor ANOVA with Fisher’s Least Significant Difference test. Statistical analyses were performed using GraphPad Prism 8 (GraphPad Software, La Jolla, California, USA) or the R language programming environment.

## DATA AND CODE AVAILABILITY

The datasets generated and analysed in this study are included in the manuscript. The raw fastq files for all of the sequencing data are available upon request and will be uploaded to GEO upon publication. Descriptions of the analyses, tools and algorithms are provided in the methods section of this article. Custom code for generating gRNA counts from fastq files and code for generating qGI-scores will be made available on Github upon publication.

## SUPPLEMENTAL FIGURE LEGENDS

**Figure S1.**
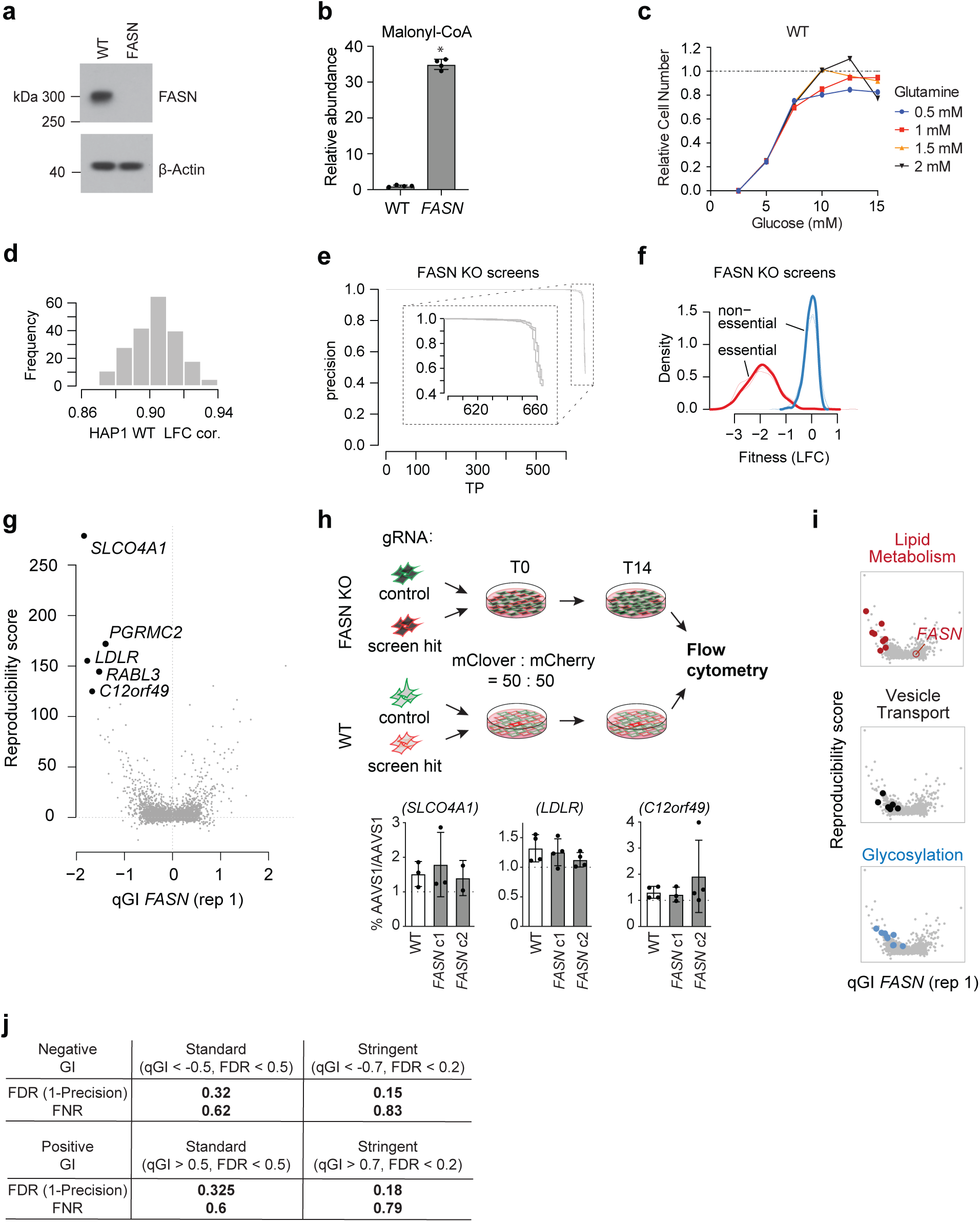
Validation of *FASN*-KO cells and genetic interactions screens. (**a**) Western blot depicting FASN and β-Actin levels in HAP1 parental wildtype (WT) and *FASN*-KO cells. (**b**) Bar plot depicting malonyl-CoA levels in HAP1 WT and *FASN*-KO cells as detected by mass spectrometry-based metabolite profiling, normalized to parent HAP1 WT cells (n=4); p = 0.03, Mann Whitney U test. (**c**) Growth curves of HAP1 WT cells depicting relative cell numbers over 3 days, plotted as a function of glucose concentration in mM, in either 0.5 mM (blue), 1 mM (red), 1.5 mM (yellow), or 2 mM (black) glutamine. (**d**) Histogram showing a frequency distribution of all pairwise Pearson correlation coefficients for LFC values (T0/T18) of the 21 WT HAP1 screens. (**e**) Precision-recall curves for the three CRISPR replicate screens in HAP1 *FASN*-KO cells using the reference core essential gene set (CEG2) defined in Hart *et al*., 2017. (**f**) Fitness effect (log2 fold-change, LFC) distributions for reference core essential (CEG2) and non-essential gene sets defined in Hart *et al*., 2017 across the three *FASN*-KO query screens. (**g**) Scatter plot showing reproducibility scores as a function of qGI scores for a single FASN-KO screen (replicate A). Pairwise reproducibility of a qGI score was calculated by computing each gene’s contribution to the covariance between a pair of screens divided by the sum of standard deviations. The reproducibility score represents the sum of those values across the three pairwise comparisons. Five genes with highest reproducibility scores and the most negative qGI scores with the *FASN*-KO screen (replicate A) are labelled. (**h**) Establishing the *AAVS1* target locus as a good negative control site in HAP1 WT and *FASN*-KO cells. Schematic depicting co-culture validation assays (upper panel). Parental WT and *FASN*-KO cells were stably transduced with color-coded gRNA expression vectors carrying an intergenic control or screen hit-targeting gRNA. Color-coded cells are mixed at an equal ratio, cultured over two weeks and the relative proportion of green and red cells was quantified by flow cytometry. Control co-culture experiments performed in parallel to the validation of hit genes depicted in main Figure 1e as indicated above each barplot (lower panel). Bar plots are depicting the color ratio of cells carrying two colour-coded gRNAs targeting *AAVS1* (intergenic control) across WT and two *FASN-*KO clones as indicated. Experiments were performed with three independent gRNA targeting AAVS1 and using both color orientations. All data are represented as means ± standard deviation (n = 3 or 4). (**i**) Scatter plots reproducibility scores as a function of qGI scores for the negative genetic interaction hits depicted in Figure 1h functioning in lipid uptake and homeostasis (red), vesicle transport genes (black) and glycosylation (blue). (**j**) Precision and recall values for GIs with *FASN* measured at the standard (|qGI| > 0.5, FDR < 0.5) and stringent (|qGI| > 0.7, FDR < 0.2) thresholds. Precision and recall values were computed using an MCMC-based approach (see Methods).

**Figure S2.**
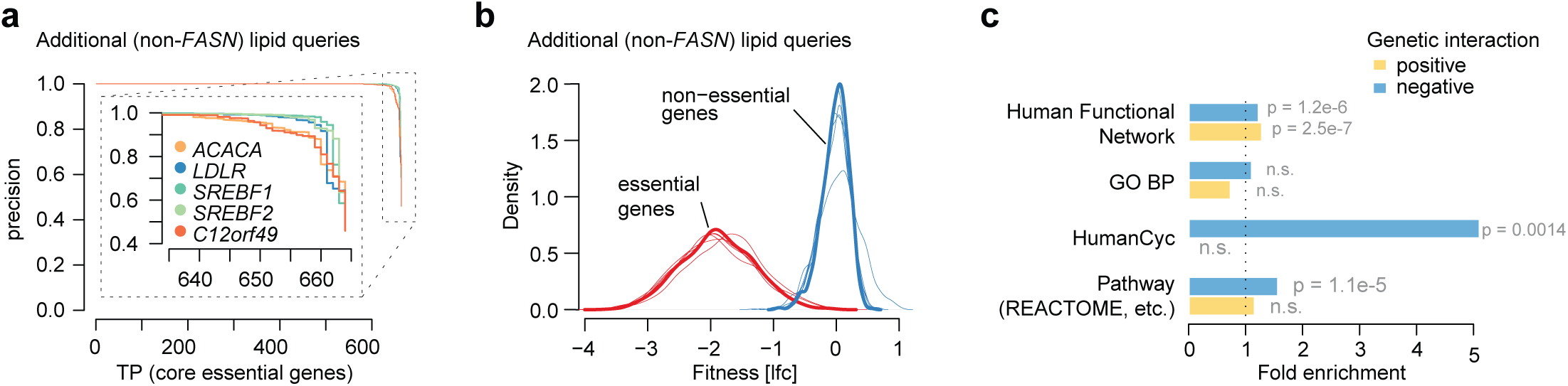
Quality control of genetic interaction screens for fatty acid synthesis-related query genes. (**a**) Precision-recall curves distinguishing the reference core essential gene set (CEG2) defined in Hart *et al*., 2017 and a non-essential gene set in CRISPR screens in five HAP1 knockout query cell lines (*LDLR, C12orf49, SREBF2, ACACA, SREBF1*-KO). (**b**) Fitness effect (LFC) distributions for reference core essential (CEG2) and non-essential gene sets defined in Hart *et al*., 2017 across CRISPR screens in five HAP1 KO cell lines (*LDLR, C12orf49, SREBF2, ACACA, SREBF1*). (**c**) Bar plot of enrichment of co-annotation as defined by the Human Functional Network, Gene Ontology Bioprocesses (BP), HumanCyc or and aggregation of pathway standards (including REACTOME, KEGG or BIOCARTA) for genetic interactions identified across all six query genome-wide screens (*FASN, LDLR, C12orf49, SREBF2, ACACA, SREBF1*). See methods for details of analysis.

**Figure S3.**
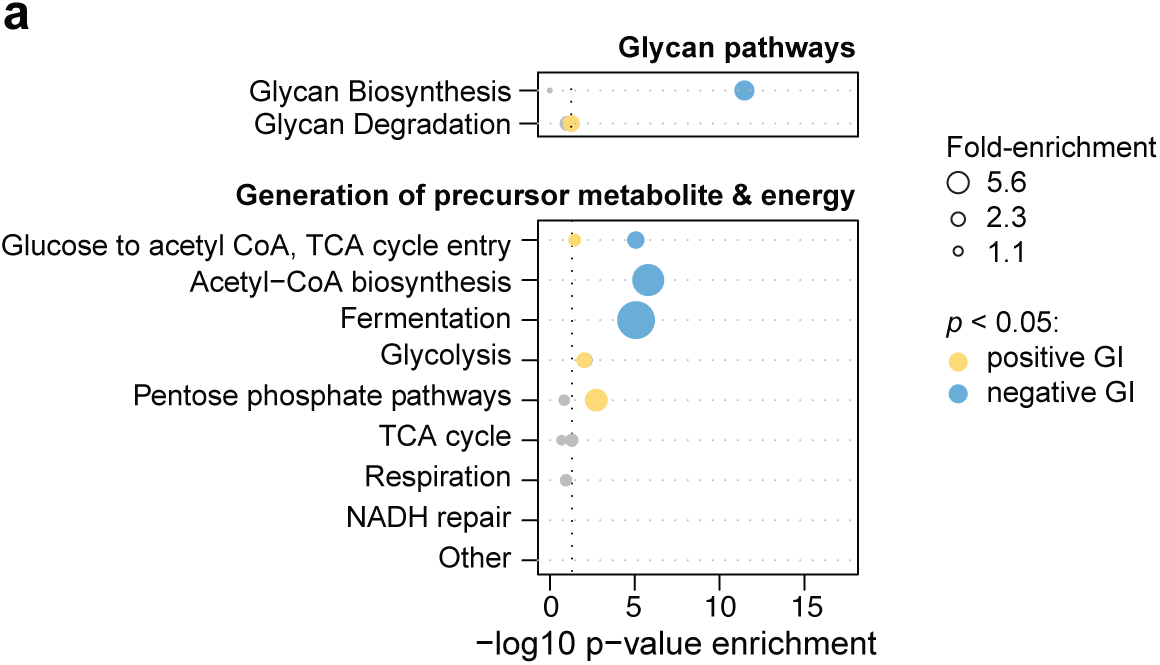
Pathway enrichment analysis of genetic interactions for fatty acid synthesis-related query genes in additional HumanCyc sub-categories. (**a**) Dot plot of normalized pathway enrichment values for aggregate GIs across the six query genes (*FASN, C12orf49, LDLR, SREBF2, ACACA, SREBF1)* with sub-categories from HumanCyc are indicated. A GI is identified for a query-library pair if the |qGI| > 0.5 and FDR < 0.5. Enrichment for positive (yellow) and negative (blue) GI is tested inside Glycan Pathways and Generation of precursor metabolite and energy HumanCyc branches using a hypergeometric test. Enrichment with p-value < 0.05 are blue (negative GI) and yellow (positive GI). Dot size is proportional to the fold-enrichment in the indicated categories and specified in the legend.

**Figure S4.**
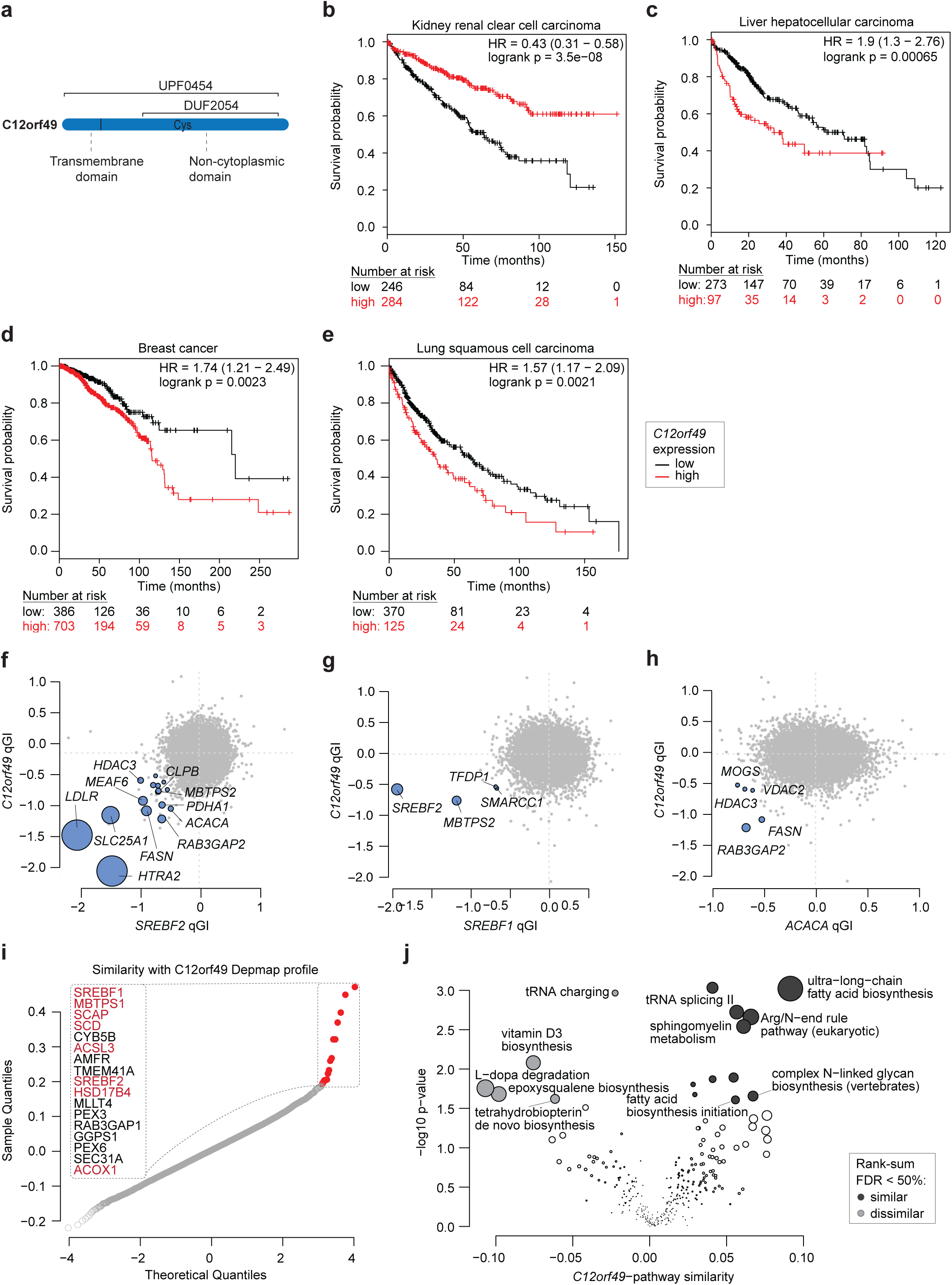
Overview of C12orf49, cancer associations, and functional correlates. (**a**) Cartoon of C12orf49 protein sequence features and domains. (**b-e**) Kaplan Meier survival plots displaying univariate analysis of TCGA data across multiple tumor types including kidney, breast, liver and sarcoma for *C12orf49* high vs. low expressing tumor tissue (www.kmplot.com) (Nagy *et al*, 2018). (**f-h**) GI overlap between *C12orf49* and *SREBF2, SREBF1* and *ACACA* qGI scores shown as pairwise scatter plots with *C12orf49* as function of *SREBF2* (f), *SREBF1* (g) and *ACACA* (h). A common negative GI is called if it is significant (qGI < −0.5, FDR < 0.5) in both screens (indicated in blue). The top 10 strongest common GIs and lipid metabolism genes are labelled. (**i**) Profile similarity of *C12orf49* across genome-wide DepMap CRISPR/Cas9 screens. Similarity was quantified by taking all pairwise gene-gene Pearson correlation coefficients of CERES score profiles across 563 screens (19Q2 DepMap data release). The distribution of 17,633 CERES profile similarity is plotted as a quantile-quantile plot, and the top 18 most similar out of 17,633 genes are labelled. Genes associated with lipid metabolism are indicated in red. (**j**) Pathway analysis of *C12orf49* profile similarity. *C12orf49* profile similarity scores for all 17,634 genes represented in the DepMap were mean-summarized by pathway as defined in the HumanCyc standard (Romero *et al.*, 2004). Tendencies towards pathway-level similarity (co-essentiality) and dissimilarity (exclusivity) with *C12orf49* were tested using a two-sided Wilcoxon rank-sum test with multiple hypothesis correction using the Benjamini and Hochberg procedure.

**Figure S5.**
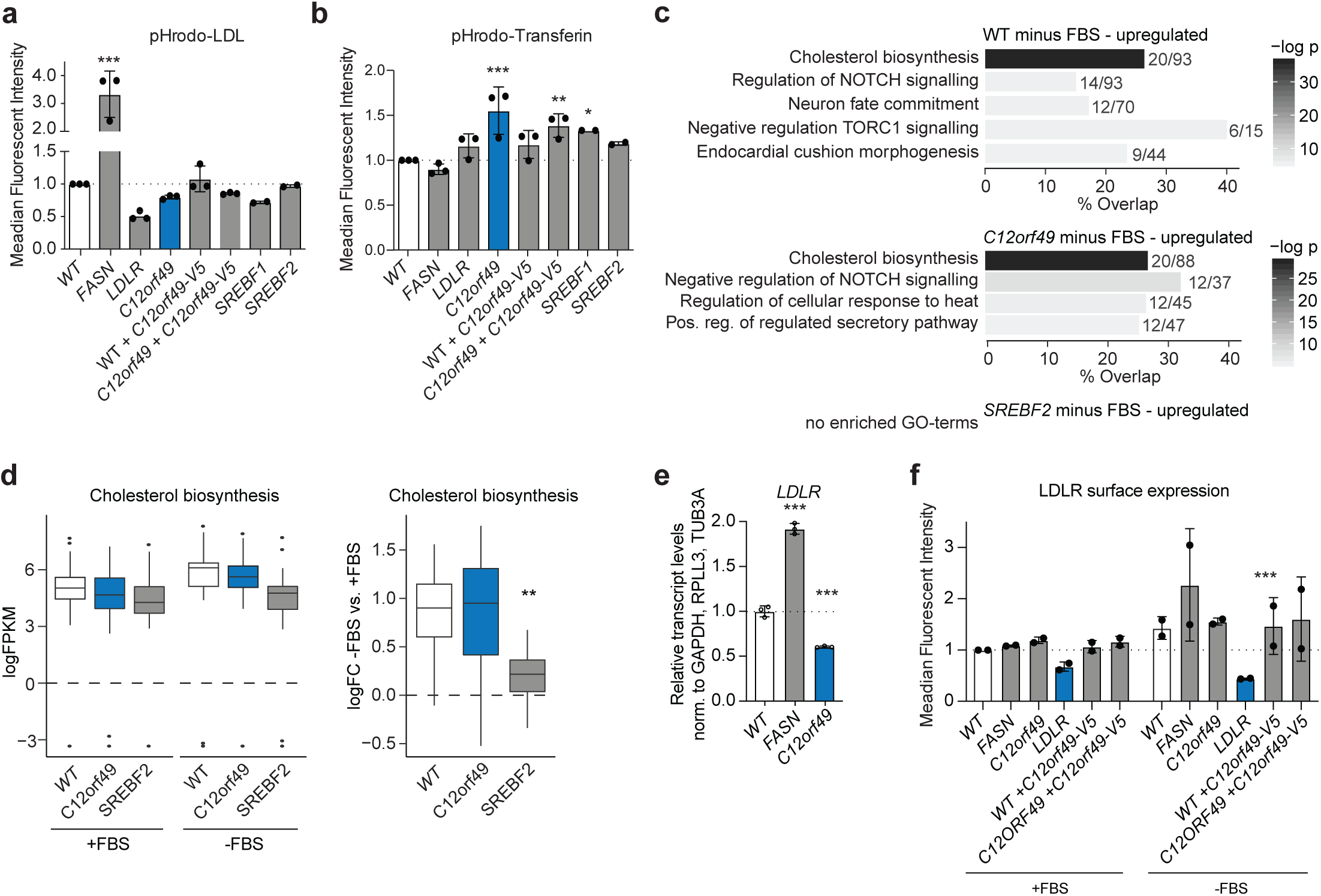
Regulation of LDL uptake and LDLR expression by *C12orf49*. (**a**) Bar plots showing the results of a low density lipoprotein (LDL) uptake assay across the indicated HAP1 cell lines using pHrodo-LDL probe. All data are represented as means ± standard deviation (n = 2-3), ***p < 0.001, one-way ANOVA. (**b**) Bar plots showing the results of a transferin uptake assay across the indicated HAP1 cell lines using pHrodo-Transferin probe. All data are represented as means ± standard deviation (n = 2-3). ***p < 0.001, **p < 0.01, and *p < 0.05; one-way ANOVA. (**c**) Gene ontology enrichment analysis of genes upregulated under serum starvation in HAP1 wildtype (WT), *C12orf49* or *SREBF2-*KO cells using GO molecular functions, GO bioprocesses and Reactome standards. Gene sets with overlapping members have been merged and bars depict mean percentage overlap with the indicated term. Numbers indicate the mean overlap and term sizes respectively. (**d**) Boxplots depicting mean expression and induction of genes assigned with the indicated term across HAP1 WT, *C12orf49* and *SREBF2-*KO cells under normal (+FBS) and serum-starved (-FBS) conditions (n=3), **p < 0.01, student’s t test. (**e**) Bar plot of relative mRNA expression of LDLR across HAP1 WT, *FASN*-KO and *C12orf49*-KO cells (n=3). ***p < 0.001, one-way ANOVA. (**f**) Bar plot of LDLR surface expression across HAP1 wildtype (WT) and the indicated KO and rescue cell lines under normal (+FBS) or serum-starved (-FBS) conditions as assessed by flow cytometry. ***p < 0.001, two-way ANOVA.

## SUPPLEMENTAL TABLES

Table S1 FASN qGI reproducibility analysis across *FASN* replicate screens.

Table S2 Genetic Interaction Scores.

Table S3 Pathway Enrichment Negative Genetic Interactions *FASN*, *C12orf49*

Table S4 Pathway enrichment Genetic interactions z-scores.

Table S5 BioID C12orf49.

Table S6 Pathway enrichment BioID C12orf49.

Table S7 RNAseq HAP1 WT, *C12orf49*-KO, *SREBF2*-KO plus-minus FBS.

Table S8 Primer and Oligo List.

